# Multiomic dissection of HR^+^/HER2^−^ invasive lobular breast carcinoma reveals mobilized yet dysfunctional anti-tumor immunity shaped by tumor-stroma crosstalk and impaired antigen presentation

**DOI:** 10.64898/2026.05.28.728418

**Authors:** Maelle Picard, Pascal Finetti, Arnaud Guille, Gwenaël Lumet, Lenaïg Mescam, Laurys Boudin, Anthony Goncalves, François Bertucci, Emilie Mamessier

## Abstract

**Context:** Immunotherapy based on immune checkpoint inhibitors (ICI) revolutionized the treatment of triple-negative (TN) breast carcinomas (BC), but remains more challenging in HR^+^/HER2^-^ BCs. Because invasive lobular carcinomas (ILC) generally exhibit low immune infiltration, ICIs were largely overlooked in this pathological type. The only clinical trial of ICIs dedicated to ILCs showed disappointing results, notably in HR^+^/HER2^-^ cases. The immune landscape of HR^+^/HER2^-^ ILCs has been poorly described. High level of tumor-infiltrating lymphocytes (TIL) was associated with worse prognosis in HR^+^/HER2^-^ ILCs. A better characterization of the immune landscape of HR^+^/HER2^-^ ILCs could clarify the poor efficiency of ICIs and the negative prognostic value of TILs, and reveal complementary targets able to increase immunotherapy efficiency.

**Method:** We comprehensively characterized the immune landscape of HR^+^/HER2^-^ ILCs, comparatively to HR^+^/HER2^-^ invasive ductal carcinomas (IDC), by applying multi-omics and multi-scale analysis (gene expression at the bulk and single-cell levels, and protein-based spatial analysis) to clinical samples.

**Results:** While the overall level of immune infiltration was comparable between both pathological types, the quality of immune infiltrate differed markedly. Comparatively to HR^+^/HER2^-^ IDCs, HR^+^/HER2^-^ ILCs were enriched in immune cells and tertiary lymphoid structures with anti-tumor potential, presented more spatial proximity between cancer cells and CD8^+^ cytotoxic T cells, and stronger theorical vulnerability to ICIs. However, in HR^+^/HER2^-^ ILCs, anti-tumor response was defective; CD8^+^ cytotoxic T cells failed to fully unleash their cytotoxic function and CD4^+^ helper T cells evidenced a pro-tumoral and naive phenotype. Furthermore, antigen-presenting compartment was defective, altogether embedded in a stronger immunosuppressive environment, enriched in immunoregulatory cancer-associated fibroblasts (iCAF).

**Conclusion:** This study contributes to explain the lesser efficiency of PD-1/PD-L1-based ICIs in HR^+^/HER2^-^ILCs by comparison with HR^+^/HER2^-^ IDCs, by shedding light on a complex ecosystem where tumor cells shape a distinctive stroma that contribute to prevent anti-tumor immune response activation. Altogether, our findings further support the rationale for combining iCAF-targeting strategy with an *ad hoc* immunotherapy (such as an anti-VTCN1/B7-H4 antibody-drug conjugates for example).

**Graphical abstract:** 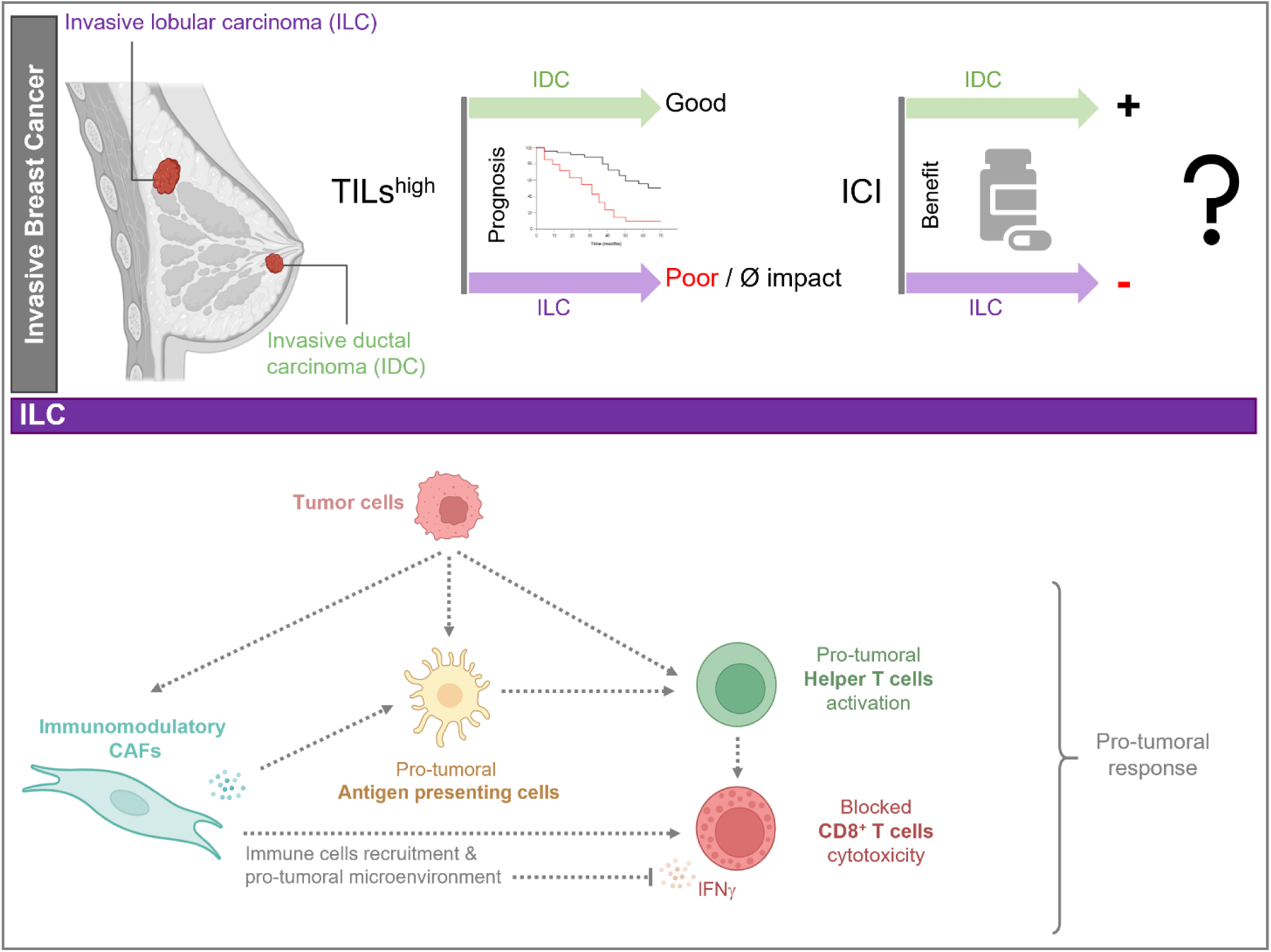

**Highlights:** ***WHAT IS ALREADY KNOWN ON THIS TOPIC**:*

- Immune cells infiltrate both HR^+^/HER2^-^ IDC and HR^+^/HER2^-^ ILC tumors, but current ICIs are less effective in HR^+^/HER2^-^ ILCs than HR^+^/HER2^-^ IDCs.

*WHAT THIS STUDY ADDS:*

- The anti-tumor immune response is mobilized but not effective in HR^+^/HER2^-^ ILCs.
- A complex ecosystem - composed of immunoregulatory cancer-associated fibroblasts, high levels of TGFâ, prostaglandin, acidosis, and a lack of antigen-presenting cells - prevents anti-tumor CD8^+^ cytotoxic T cell activation in HR^+^/HER2^-^ ILCs.

*HOW THIS STUDY MIGHT AFFECT RESEARCH, PRACTICE, OR POLICY:*

- Targeting the PD-1/PD-L1 axis is not the appropriate therapeutic strategy for HR^+^/HER2^-^ ILCs. A more complex approach should be considered, notably those combining other immune-based strategies and iCAF targeting, which may offer a better chance to eradicate HR^+^/HER2^-^ ILC tumor cells.

## Introduction

Invasive lobular carcinoma (ILC) is the second most common pathological type of breast cancer (BC) after invasive ductal carcinoma (IDC), also known as invasive BC of no special type. It accounts for approximately 10 to 15% of all diagnosed invasive BCs, a significant proportion given the high incidence of BC worldwide. Although often diagnosed in older patients, the incidence of ILCs is increasing in women aged of 45 to 50 years, possibly due to the widespread use of hormone replacement therapy ^1^ ^2^ ^3^

ILCs are characterized by small, round, and poorly cohesive neoplastic cells that invade the stroma in a single-file pattern and form indistinct margins. The poorly cohesive nature is due to loss of the *CDH1* gene (loss of 16q, the hallmark of ILCs), which encodes the E-cadherin protein, in 90% of ILC cases ^4^. Most ILCs are hormone receptor-positive (HR^+^), *i.e.* estrogen receptor-positive (ER^+^) and/or progesterone receptor-positive (PR^+^), but HER2-negative (HER2^-^) ^3^; they have a low to intermediate mitotic index, and a low pathological grade, indicating that ILCs are predominantly of the Luminal A subtype. All of these features are generally considered as good-prognosis factors ^4^ ^5^ ^6^. However, ILCs remain difficult to diagnose, as the lesions are barely palpable and hard to detect with positron emission tomography scans and mammography ^3^. Compared to IDCs, HR^+^/HER2^-^ ILCs are more often bilateral at diagnosis and three times more likely to metastasize to the peritoneum, gastrointestinal tract and ovaries ^7^. Patients with metastatic ILCs display shorter overall survival than patients with HR^+^/HER2^-^ metastatic IDCs ^8^.

The standard systemic treatment for HR^+^/HER2^-^ ILCs includes chemotherapy, hormone therapy, and CDK4/6 inhibitors. Developing new therapies is crucial for ILCs ^9^. However, patients are often underrepresented in clinical trials due to the lack of measurable metastatic disease according to RECIST, and subgroup analyses dedicated to ILCs are very rare ^10^. Regarding immunotherapy based on immune checkpoint inhibitors (ICIs), it achieved notable success in the more immunogenic triple-negative (TN) BC, but its application in HR^+^/HER2^-^ BC, traditionally classified as immunologically “cold”, remains more challenging. Recently, randomized clinical trials indicated that addition of ICI to neoadjuvant chemotherapy significantly improved the rate of pathological complete response in selected high-risk HR^+^/HER2^-^ patients ^11^ ^12^. In these trials, patients with ILCs were excluded.

Indeed, ILCs have long been considered weakly immunogenic tumors due to a low percentage (∼5%) of tumor-infiltrating lymphocytes (TIL). However, the recent characterization of immune infiltrate in ILCs suggested potential interest in using ICIs in specific cases ^6^ ^13^. Indeed, profiling of ILCs identified different molecular subtypes: the *hormone-related* subtype, and the *immune-related* subtype ^14^ or, in another study, the *reactive-like* subtype, the *immune-related* subtype and the *proliferative* subtype ^4^. The immune-related subtype was robust enough to be distinguished in both studies. Based on these observations and the clinical partial responses reported in two of three HR^+^/HER2^-^ ILC patients treated with ICIs in the KEYNOTE-028 trial (NCT02054806, ^15^), the phase II GELATO trial (NCT03147040, ^16^) recently tested the combination of weekly carboplatin and atezolizumab (PD-L1 inhibitor) specifically in metastatic ILC patients ^16^. Four of twenty-three evaluable patients had a partial response (17%), and two had stable disease; four of these six patients had TN ILCs. Patients with TN ILCs had a significantly higher clinical benefit rate than those with HR^+^/HER2^-^ILCs. As responses were short-lived and observed mainly in TN ILCs, the trial stopped accrual after the first stage was completed.

This disappointing result in patients with HR^+^/HER2^-^ ILCs, contrasting with existence of an immune-related subtype, suggests that a more nuanced approach to immunotherapy is required to significantly impact outcomes for HR^+^/HER2^-^ ILC patients. However, it is interesting to note that high TILs infiltrate was associated with a worse prognosis in two independent ILC cohorts ^17^. Thus, to make progress, a better understanding of immune cellular composition and functional states in ILC tumors is crucial.

In this line, a recent study shed light on a mechanism that may contribute to immunotherapy failure in HR^+^ ILCs using integrative analysis of single-cell RNA sequencing (scRNAseq) and multispectral immunohistochemistry. The macrophage landscape varied between HR^+^ IDCs and HR^+^ ILCs, with those in HR^+^ IDCs displaying predominantly anti-tumor features, whereas those in HR^+^ ILCs exhibited a pro-tumor phenotype. These findings suggest that macrophage-repolarizing immunotherapies could provide clinical benefit in HR^+^ ILCs to overcome resistance to current approaches. The study also considered two hypotheses that were not fully explored: first, that immunotherapy failure may stem from improper or suboptimal T cell priming and activation, consistent with low dendritic cell infiltration; and second, that the enrichment of regulatory T cells and myeloid-derived suppressor cells within the tumor microenvironment of HR^+^ ILCs could further dampen therapeutic efficacy ^18^.

Because the tumor microenvironment has multiple ways of impairing T-cell activation, we used multi-omics analysis, complemented with multi-scale approaches, to comprehensively characterize the tumor immune infiltrate and microenvironment of HR^+^/HER2^-^ ILCs comparatively to those of HR^+^/HER2^-^ IDCs. We revealed details on the mobilization of the anti-tumor immune compartment and its embedment in a more immunosuppressive tumor microenvironment compared to IDCs. The complex ecosystem we described in HR^+^/HER2^-^ ILCs may provide a preclinical basis for understanding their poor sensitivity to ICIs and identifying complementary therapeutic targets.

## Materials and Methods

The general workflow of this study was summed up in Figure 1A.

**Figure 1:**
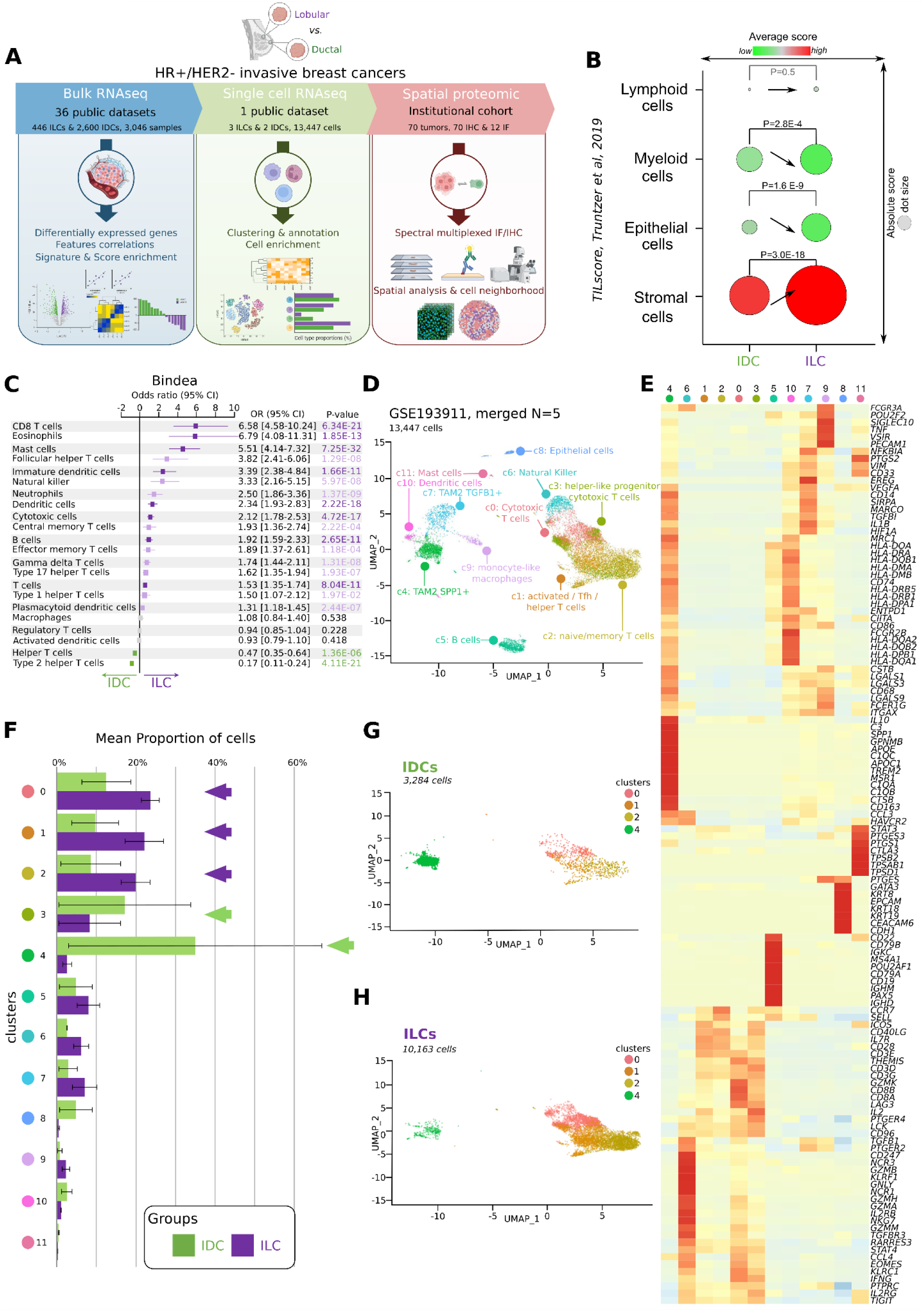
Immune infiltrate was similar in quantity but differed in quality between HR^+^/HER2^-^ ILCs and HR^+^/HER2^-^ IDCs. (A) General workflow of the study. RNAseq; RNA sequencing, IF; immunofluorescence, IHC; immuno-histochemistry, HR; hormone receptor, ILC; invasive lobular carcinoma, IDC; invasive ductal carci-noma. Created in BioRender. Mamessier, E. (2026) https://BioRender.com/ x3x0w33 (B) Dot plot representing the enrichment in lymphoids, myeloids, epithelial cells and stroma TILscore (Truntzer *et al*) in HR^+^/HER2^-^ ILCs *vs* HR^+^/HER2^-^ IDCs from bulk RNA sequencing datasets. The dot size represented the absolute score and the green-to-red scale represents the relative average score. Arrows summary the enrichment between HR^+^/HER2^-^ ILCs and HR^+^/HER2^-^ IDCs; flat means nearly no differences, increasing means enrichment in HR^+^/HER2^-^ ILCs and decreasing means loss in HR^+^/HER2^-^ ILCs. (C) Tree plot representing the enrichment in Bindea *et al* immune signatures in HR^+^/HER2^-^ ILCs *vs* HR^+^/HER2^-^ IDCs from bulk RNA sequencing datasets. CI; confidence interval. P-values in dark purple referred to most significative p-value (P> 1E-10), then light purple ones to mid significative p-value (5E-2<P<1E-10) in HR^+^/HER2^-^ ILCs whereas green ones referred to significant p-value enriched in HR^+^/HER2^-^ IDCs. Grey frames are for readability. Green is for IDCs and purple is for ILCs and this color code will be used throughout the article. (D) Uniform Manifold Approximation and Projection (uMAP) representing the clustering of the 13,447 cells recovered from the five patients of the scRNAseq GSE193911 dataset (Onkar *et al*) and their annotation. Tfh; T follicular helper, TAM2; type 2 tumor-associated macrophages, c; cluster, N; ef-fective. (E) Hierarchical heatmap representing the differentially expressed genes used to annotate clusters ob-tained from the scRNAseq GSE193911 (Onkar *et al*) datasets (Figure 1D). Clusters colors were re-minded for readability. (F) Barplot representing the cell proportion of each cluster in HR^+^/HER2^-^ ILCs *vs* HR^+^/HER2^-^ IDCs pa-tients. Arrows indicated clusters with main difference between cluster’s proportions in HR^+^/HER2^-^ILCs *vs* HR^+^/HER2^-^ IDCs. Clusters colors were reminded for readability. (G) uMAP representing the clustering of the four clusters of interest (Immune_c4, Immune_c2, Im-mune_c1 and Immune_c0) in HR^+^/HER2^-^ IDCs patients. Clusters colors were reminded for readabil-ity. (H) uMAP representing the clustering of the four clusters of interest (Immune_c4, Immune_c2, Im-mune_c1 and Immune_c0) in HR^+^/HER2^-^ ILCs patients. Clusters colors were reminded for readabil-ity.

### Gene expression analysis of bulk tumor samples

#### Publicly available gene expression database

We retrospectively analyzed clinicopathological and gene expression data of bulk clinical samples from 36 public data sets ^19^, including ours ^20^. They had been profiled using DNA microarrays or RNA sequencing and data were gathered from the National Center for Biotechnology Information (NCBI)/Genbank GEO, ArrayExpress, European Genome-Phenome Archive, The Cancer Genome Atlas portal (TCGA) databases, and authors’ website. The details of institutional review board and ethical committee approval and patients’ consent for all studies are present in their corresponding publications listed in Supplementary Table 1. The series comprised a total of 8,982 non-redundant primary BC samples.

#### Analysis of public gene expression data

Several steps of pre-analytic data processing were applied as previously described ^21^. Briefly, each set was first separately normalized according to its platform: Robust Multichip Average with the non-parametric quantile algorithm for the raw Affymetrix data, and quantile normalization for the already processed data sets (Agilent, SweGene, Illumina microarrays and RNA sequencing log2-transformed data). Then, expression levels of genes of interest were standardized within each data set using the PAM50 Luminal A population as reference. This allowed us to exclude biases due to laboratory-specific variations and to population heterogeneity and to make data comparable across all sets. To avoid biases related to trans-institutional immunohistochemistry analysis and thanks to the bimodal distribution of respective mRNA expression levels, the ER, PR, and HER2 statutes (negative/positive) were defined from transcriptional data of *ESR1*, *PGR*, and *HER2* respectively, as described ^22^. The molecular subtypes of samples were then defined as HR^+^/HER2^-^ for ER-positive and/or PR-positive and HER2-negative tumors, HER2^+^ for HER2-positive tumors, and TN for ER-negative, PR-negative, and HER2-negative tumors. Of the 8,982 primary tumor samples, we identified 3,046 HR^+^/HER2^-^ tumor samples including 446 ILCs and 2,600 IDCs. Next, we applied several multigene signatures to each normalized data set separately. They were selected because of their relevance in BC and in immunity, and included: the PAM50 molecular subtypes ^23^; prognostic signatures such as Recurrence Score ^24^ and 70-gene signature ^25^; immune-related signatures such as TILs score ^26^, signatures of 24 different innate and adaptive immune cell subpopulations ^27^, CIBERSORT scores ^28^, Tertiary Lymphoid Structure (TLS) score ^29^, T cell inflammation signature (TIS) score ^30^, Immunologic Constant of Rejection (ICR) ^19^, Cytolytic activity score ^31^, LCK signature ^32^, and Gatza’s pathway activation scores ^33^.

We also conducted a supervised analysis of expression profiles of 1,900 tumor samples (learning set) to search for differentially expressed genes (DEG) between the 317 HR^+^/HER2^-^ ILCs and the 1,583 HR^+^/HER2^-^IDCs. This learning set included the two largest transcriptomic datasets (TCGA and Metabric), which were merged and batch-corrected using ComBat algorithm (sva R package). The effectiveness of batch correction was assessed through Principal Component Analysis (PCA) (Supplementary Figure 1A). Supervised analysis was performed using a moderated t-test with empirical Bayes statistic included in the limma R packages. False discovery rate ^34^ was applied to correct the multiple testing-hypothesis. Significant genes were defined by p<5%, q<10%, and fold change superior to |1.5x|. These genes were used to construct a centroid as the HR^+^/HER2^-^ ILCs median expression profile within the learning set and, for each sample, a Pearson correlation score was calculated as distance parameter. This score was then compared between the HR^+^/HER2^-^ ILCs and HR^+^/HER2^-^ IDCs samples. Robustness of the resulting gene list was tested in the validation set of 1,146 remaining HR^+^/HER2^-^ BC samples (129 HR^+^/HER2^-^ ILCs and 1017 HR^+^/HER2^-^ IDCs). Ontology analysis of the resulting gene list was based on the Database for Annotation, Visualization and Integrated Discovery (DAVID; http://david.abcc.ncifcrf.gov/). Data processing was done in R using Bioconductor and associated packages (R-4.1.2, Bioconductor-1.30.21.1).

#### Statistical analysis

Correlations between tumor classes and clinicopathological variables were analyzed using the Student t-test or Fisher’s exact test when appropriate. Overall survival (OS) was calculated from the date of diagnosis until the date of cancer-related death, and disease-free survival (DFS) was calculated from the date of diagnosis until the date of relapse (local, regional, or distant) or death from any cause. Follow-up was measured from the date of diagnosis to the date of last news for event-free patients. Survivals were calculated using the Kaplan-Meier method and curves compared with the log-rank test. All statistical tests were two-sided at the 5% level of significance. Statistical analysis was done using the survival package (version 3.5-5) in the R software (version 4.1.2; http://www.cran.r-project.org/).

#### Single-cell RNA sequencing

ScRNAseq data were obtained from a publicly available dataset (GSE193911, ^18^). From this dataset, we collected data corresponding to five patients with HR^+^ BCs (3 ILCs and 2 IDCs) for which TILs were available and not multiplexed with peripheral blood mononuclear cells. A complementary dataset consisting of 1 ILC and 10 IDCs for which epithelial tumor cells were available was collected (EMTAB10607,^35^) to precise the expression of *VTCN1/B7-H4*. ScRNAseq analysis was performed using the R package Seurat v4.3.0.1. The five samples were merged into a single dataset. Cells with less than 350 genes *per* cell and less than 180 unique molecular identifiers (UMI) *per* cell were filtered out. Subsequently, cells with more than 25,000 UMIs *per* cell and a mitochondrial percentage exceeding 20% were removed. This resulted in a final dataset of 13,447 filtered cells. Data normalization and scaling were performed, followed by cell cycle score regression. PCA was computed using 50 principal components, and Harmony was employed to correct for batch effects. Louvain clustering with a resolution parameter of 0.8 was used to identify 11 clusters, differing from the original study. Finally, Uniform Manifold Approximation and Projection (uMAP) was used to visualize the 11 clusters. Cell subtypes were annotated in regards to the presence of markers among the DEG. Signature score enrichment was computed using Ucell package. Pseudo-bulk DEG analysis was performed for each cluster between HR^+^/HER2^-^ ILCs and HR^+^/HER2^-^ IDCs. Genes were considered differentially expressed with an adjusted p-value threshold of 5% and a log2 fold-change threshold of 0.58.

#### Patients and breast cancer samples for protein expression analyses

Protein expression analyses were applied to a retrospective series of samples collected from patients with primary ILCs or IDCs treated in our institution. Main inclusion criteria were: female adult patients diagnosed from 2015, with a histologically proven non-inflammatory and non-metastatic ILCs or IDCs; diagnosis confirmed by a BCs expert pathologist; primarily treated by surgery; with formalin buffer and paraffin-embedded (FFPE) sample; with informed status for HR (ER and PR both positive) and HER2 (negative); treated by hormone therapy at the Paoli-Calmettes Institute; and with informed consent available. Clinical features, treatment modalities and outcomes were obtained from the medical records. A total of 34 HR^+^/HER2^-^ ILCs, and 36 HR^+^/HER2^-^ IDCs (comparative controls) were obtained from pathology department archives. The study was approved by our institutional review board (ILC_HR+-IPC 2022-010). All 70 samples were used for immunohistochemistry (IHC) and 12 for immunofluorescence (IF). Sample preparation, staining and read-out were performed in our Predictive Oncology laboratory. Their clinicopathological characteristics are summarized in Supplementary Table 2.

### Multiplexed immunohistochemistry

#### Staining

Tumor samples from 70 patients (34 HR^+^/HER2^-^ ILCs and 36 HR^+^/HER2^-^ IDCs) were analyzed. Three-µm tissue sections were cut with a microtome (Epredia™ HM 340E Electronic Rotary Microtome, Fisher Scientific, USA) and mounted on TOMO® Adhesion Microscope Slides (Matsunami GlassSlides, Japan). Slides were dried at room temperature for 30 minutes before an overnight dewaxing at 56°C. Tissues underwent an epitope retrieval (pH=8.0) for 8 minutes. Staining was performed following a personalized Discovery Ultra procedure; briefly, CD20 primary antibody (NCL-L-CD20-L26, Leica, Germany) was incubated for 1 hour at room temperature, anti-mouse secondary antibody was incubated for 16 minutes at room temperature followed by a DAB staining for 8 minutes. Tissues underwent an epitope retrieval (pH=6.0) for 8 minutes allowing to removed antibodies in excess. CD3 primary antibody (GA503, DAKO Agilent, USA) was incubated for 1 hour at room temperature, anti-rabbit secondary antibody was incubated for 16 minutes at room temperature, followed by a purple staining for 36 minutes. Finally, a counterstaining was performed using hematoxylin for 8 minutes and Bluing reagent for 4 minutes. Slides were mounted using Dako Coverslipper (Agilent, USA).

#### Image acquisition and processing

Whole-slide images were acquired using NanoZoomer 2.0-HT™ (Hamamatsu Photonics, Japan) and processed using QuPath-0.5.1. CD3^+^ and CD20^+^ lymphocytes were detected using positive cell detection tools with suitable settings for cell size, nuclei counterstaining signal and positive labeling signal. Then, we used the density map tool to identify aggregates of T and/or B cells (more than 100 cell clusters). The number of lymphoid aggregates *per* tissue was counted and their architectural organization was sorted by a pathologist into “unorganized” (*i.e.,* with T and B cells showing a diffuse pattern), “TLS-like, immature” (*i.e.,* with T and B cells being organized with a B-cell zone, surrounded by a T-cell zone), “TLS-like, mature” (*i.e.,* with the B cells zone, showing larger, proliferating and dying B cells forming a germinal center-like zone). The total number of TLS-like structures (matures or immatures) was compared between the HR^+^/HER2^-^ ILC and HR^+^/HER2^-^ IDC groups. We considered tissues with more than two TLS-like to be non-random and indicative of a biological purpose; we also compared the frequency of tissues with more than two TLS-like structures between the two groups using a contingency test.

### Multiplexed Immunofluorescence

#### Region of interest selection

We selected 12 patients (6 IDCs and 6 ILCs) with BCs presenting an immune infiltrate. Tissue sections of 3 µm were cut from FFPE blocks with a microtome (Epredia™ HM 340E Electronic Rotary Microtome, Fisher Scientific, USA) and mounted on blue slides Klinipath™ (Avantor, USA). Slides were dried at room temperature before being colored by Hematoxylin-Eosin-Safran. Slides were scanned using the NanoZoomer 2.0-HT™ (Hamamatsu Photonics, Japan) and 3 to 10 regions of interest (ROI) *per* tissue were selected using the CaloPix software (version 2.10.16) when available.

#### Staining

Serial 3-µm tissue sections were cut with a microtome (Epredia™ HM 340E Electronic Rotary Microtome, Fisher Scientific, USA) and mounted on TOMO® Adhesion Microscope Slides (Matsunami GlassSlides, Japan), dried at room temperature before being incubated overnight at 56°C for a first step of dewaxing. Slides underwent a 10-minute-long wash of Histolemon® (CARLO ERBA, Italia) to complete dewaxing, then a progressive ethanol bath to rehydrate the tissue. Slides underwent heat-induced epitope retrieval in Tris-EDTA buffer (pH=9.0) following PT Link procedure (Dako Agilent, USA).

To quench FFPE-tissue natural autofluorescence, the slides were incubated in a solution of Sudan Black B 0.3% (Sigma Aldrich, USA) ethanol 70% for 15 minutes. They were then washed with Tween20 (Euromedex, France) 0.02% phosphate buffer saline (PBS) (Gibco, Thermo Fischer Scientific, USA) 1X for 15 minutes, then PBS 1X for 5 minutes. Tissues were permeabilized with Tritonx100 (Euromedex, France) 0.5% PBS 1X for 15 minutes and saturated with Normal Horse Serum (ThermoFisher Scientific, USA) 5% PBS 1X for 30 minutes. Seven-color manual multiplexed immunofluorescence (mtpx IF) was performed using the “ImmCytScore” panel allowing to localize immune cells (CD45^+^), cytotoxic cells (CD8^+^, GZMB^+^, CD56^+^), T cells (CD3^+^) and epithelial tumor cells (pan-KRT^+^) within the tissue using manual mtpx IF, Supplementary Table 3 and SytoxBlue (1:5000 dilution in water, Miltenyi Biotech, Germany). Antibodies mix was incubated 1 hour at room temperature and obscurity. Slides were then washed several times by serial baths with saturation solution, then PBS 1X and finally with miliQ water before being slipped with Prolong antifade mountant (ThermoFisher Scientific, USA).

#### Images acquisition

A minimum of 3 to 10 ROIs (when possible) were imaged with confocal microscope (LSM800, Carl Zeiss, Germany) at x25 immersion magnification using appropriate exposure times and 2x2 tile scan and with a particular attention for the representativity of ROI acquired. Areas of interest were acquired and analyzed with an automatized software (Zen Black, version 3.3.89.0000, Carl Zeiss, Germany). Whole-slide images were acquired using NanoZoomer S60 Digital slide scanner (Hamamatsu, Japan), visualized using NDP.view 2 and processed using QuPath-0.5.1.

#### Image processing

Image files were converted from the proprietary *.czi* format to 32 bits *.tiff* image files using Zeiss software (Zen Black, version 3.3.89.0000, Carl Zeiss, Germany). Images were then visualized and annotated to identify large artefacts, *i.e.,* from hemolysis in large vessels or from large fibrous regions, using QuPath 0.5.1. Masks were generated directly in QuPath using Groovy and were used to exclude these artefacts from *.tiff* image files.

#### Single-cell segmentation and quantification

Steinbock toolkit ^36^ was used in Python (version 3.10.13) to segment the non-transformed image with the Mesmer ^37^ pre-trained neural network for single-cell segmentation. Regarding Mesmer segmentation parameters, Sytox Blue channel was used as the nuclear channel, all the remaining staining were used as membrane channels, and the pixel width was set to 0.17 µm. Signal intensities, object coordinates and objects properties were measured and exported using Scikit-image and Numpy.

#### Cell phenotyping and spatial and cellular neighborhood analysis

Single-cell properties were imported and read as a SpatialExperiment object using SpatialExperiment package in R (version 4.3.3). Intensity signals for each cell were inverse hyperbolic sine-transformed to account for the large number of zeros in image data. Transformed data along with images and mask overlays were used to perform visualization-based manual gating using EBImage (version 4.44.0) and Cytomapper (version 1.17.0) to identify and select pan-KRT^+^ cells, CD3^+^ CD8^-^ cells, CD3^+^ CD8^+^ GZMB^+^ cells, CD3^+^ CD8^+^ GZMB^-^ cells, and GZMB^+^ CD8^-^ cells.

Cell population densities were defined as the mean number of any given cell population *per* area unit *per* patient. Cell distances were computed by creating cell coordinates matrix and calculating Euclidean distances for each cell pair. To assess the proportions of CD8^+^ cells and pan-KRT^+^ with direct physical interactions, strictly heterotypic distances between CD8^+^ and pan-KRT^+^ cells were computed, and cells pairs with a distance below 25 µm were considered adjacent.

## Results

### HR^+^/HER2^-^ ILCs display more favorable prognostic features than HR^+^/HER2^-^ IDCs

We analyzed whole-transcriptomic data from 446 HR^+^/HER2^-^ ILCs and 2,600 HR^+^/HER2^-^ IDCs. Their clinicopathological and molecular features are summarized in Table 1. Compared to HR^+^/HER2^-^ IDCs, HR^+^/HER2^-^ ILCs were more frequently positive for ER and PR (96% *vs* 93%, P=2.80E-2; and 83% *vs* 75%, P=8.31E-3, respectively). They were mainly of the Luminal A subtype (44%), then Normal-like (28%) and Luminal B (20%). Compared to HR^+^/HER2^-^ IDCs, the proportion of more favorable subtypes (Luminal A and Normal-like) was higher in HR^+^/HER2^-^ ILCs than in HR^+^/HER2^-^ IDCs (44% and 28% *vs* 36% and 11%, P=6.86E-29 respectively). ILCs tumors showed higher pathological tumor size, being more often pT3 than IDCs (25% *vs* 8%, P=2.60E-26 respectively). They also displayed less grade 3 tumors than HR^+^/HER2^-^ IDCs (18% *vs* 38%, P=1.48E-8 respectively). There was an enrichment of low-risk patients in HR^+^/HER2^-^ ILCs *vs* HR^+^/HER2^-^ IDCs as defined by two key prognostic signatures: the “70-gene signature” ^25^ (P=9.77E-20) and the Recurrence Score ^24^ (P=6.79E-8). Accordingly, the incidence of DFS and OS events was lower in patients with HR^+^/HER2^-^ILCs than in those with HR^+^/HER2^-^ IDCs (17% *vs* 22%, P=3.43E-2 and 14% *vs* 19%, P=1.44E-2, respectively, Table 1).

**Table 1:** Clinicopathological characteristics of patients and samples associated with HR+/HER2- invasive lobular or ductal breast carcinomas.

| Characteristics |  | N | ILC | IDC | p-value |
| --- | --- | --- | --- | --- | --- |
| Sample |  | 3046 | 446 | 2600 |  |
| Patients' age, median, years |  | 2795 | 61 (28-90) | 60 (24-90) | 1.20E-02 |
| ER |  |  |  |  | **2.80E-02** |
|  | Negative | 196 | 18 (4%) | 178 (7%) |  |
|  | Positive | 2698 | 411 (96%) | 2287 (93%) |  |
| PR |  |  |  |  | **8.31E-03** |
|  | Negative | 332 | 44 (17%) | 288 (25%) |  |
|  | Positive | 1083 | 215 (83%) | 868 (75%) |  |
| PAM50 subtype |  |  |  |  | **6.86E-29** |
|  | Luminal A | 1120 | 195 (44%) | 925 (36%) |  |
|  | Luminal B | 1054 | 88 (20%) | 966 (37%) |  |
|  | Normal | 418 | 126 (28%) | 292 (11%) |  |
|  | Basal | 201 | 22 (5%) | 179 (7%) |  |
|  | ERBB2 | 253 | 15 (3%) | 238 (9%) |  |
| pT |  |  |  |  | **2.60E-26** |
|  | 1 | 933 | 94 (24%) | 839 (39%) |  |
|  | 2 | 1337 | 202 (51%) | 1135 (53%) |  |
|  | 3 | 268 | 100 (25%) | 168 (8%) |  |
| pN |  |  |  |  | 0.51 |
|  | 0 | 1306 | 190 (47%) | 1116 (49%) |  |
|  | 1 | 1372 | 213 (53%) | 1159 (51%) |  |
| Grade |  |  |  |  | **1.48E-08** |
|  | 1 | 248 | 38 (17%) | 210 (11%) |  |
|  | 2 | 1150 | 143 (64%) | 1007 (51%) |  |
|  | 3 | 799 | 41 (18%) | 758 (38%) |  |
| 70-gene signature |  |  |  |  | **9.77E-20** |
|  | Low risk | 969 | 225 (50%) | 744 (29%) |  |
|  | High risk | 2077 | 221 (50%) | 1856 (71%) |  |
| Recurrence Score |  |  |  |  | **6.79E-08** |
|  | Low risk | 931 | 177 (40%) | 754 (29%) |  |
|  | Intermediate risk | 893 | 142 (32%) | 751 (29%) |  |
|  | High risk | 1222 | 127 (28%) | 1095 (42%) |  |
| DFS event | N (%) | 2211 | 60 (17%) | 406 (22%) | **3.43E-02** |
| 5-year DFS | % [95%CI] | 2211 | 83% [78-88] | 85% [83-87] | 0.655 |
| OS event | N (%) | 2210 | 49 (14%) | 358 (19%) | **1.44E-02** |
| 5-year OS | % [95%CI] | 2210 | 86% [82-91] | 88% [86-90] | 0.898 |
| N, effective; ER, estrogen receptor; PR progesterone receptor; pT & pN; pathological tumor and node status; DFS, disease-free survival; OS, overall survival; CI, confidence interval; bold is for significant p-value of interest |  |  |  |  |  |

Thus, as expected ^38^ ^39^, and comparatively to HR^+^/HER2^-^ IDCs, HR^+^/HER2^-^ ILCs showed an enrichment in good-prognosis features.

### HR^+^/HER2^-^ ILCs and HR^+^/HER2^-^ IDCs have similarly low levels of immune infiltration

To study the biological pathways specific to HR^+^/HER2^-^ ILCs, we first performed a supervised analysis of gene expression profiles in our learning set (317 HR^+^/HER2^-^ ILCs *vs* 1,583 HR^+^/HER2^-^ IDCs). We identified a total of 456 DEG, including 405 overexpressed and 51 underexpressed in HR^+^/HER2^-^ ILCs *vs* HR^+^/HER2^-^ IDCs Supplementary Table 4). We confirmed the robustness of this gene list in our validation set (129 HR^+^/HER2^-^ILCs *vs* 1,017 HR^+^/HER2^-^ IDCs) (Supplementary Figure 1A-D).

Because immunotherapy based on ICIs targets the immune compartment, we investigated the degree of representation of immune genes. We first searched for canonical markers of immune cells in this signature. Focusing on the top 100 genes up-regulated in HR^+^/HER2^-^ ILC samples, only four immune-related genes where found: *IL33* that is a pro-inflammatory molecule involved in tissue repair/remodeling, *IL34* that is involved in macrophage survival and differentiation, *IL11RA* that encodes for a receptor of several cytokines and is involved in mesenchymal signaling, and *C7* a complement factor. In HR^+^/HER2^-^ IDC samples, no key gene or ontology indicated major signs of immune infiltration or activation (Supplementary Table 4, Supplementary Table 5). This result suggested that, at the transcriptional analytic level of tumor bulk, differences in immune cells are minor between both pathological types and that complementary and in-depth analyses are required to understand the involvement of the immune compartment in ICI failure in HR^+^/HER2^-^ ILCs.

To further investigate the composition of the immune infiltrates in HR^+^/HER2^-^ ILCs compared to HR^+^/HER2^-^IDCs, we first applied the TILs score, which estimates the proportions of lymphoid, myeloid and other compartments using a deconvolution strategy ^26^. Confirming our first impression and literature, the immune compartment of HR^+^/HER2^-^ BCs was not predominant and very inferior to the stromal one (Figure 1B). By comparing HR^+^/HER2^-^ ILCs and HR^+^/HER2^-^ IDCs, the lymphoid infiltrates did not show significant enrichment (P=0.5), whereas the myeloid lineage was higher in HR^+^/HER2^-^ ILCs (P=2.8E-4), as was the stromal lineage (P=3E-18). Second, we compared the prognostic value of the TILs lymphoid score in both pathological types and found that the score was associated with longer DFS and OS in HR^+^/HER2^-^ IDCs, but not in HR^+^/HER2^-^ ILCs (Table 2).

**Table 2:**
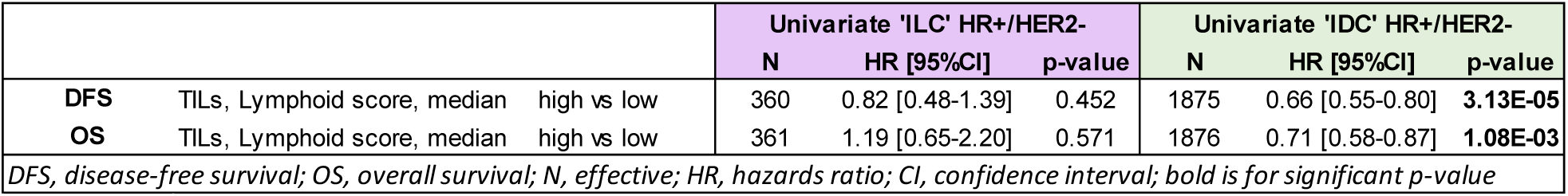
Association between prognosis and TILs lymphoid score in HR+/HER2- invasive lobular or ductal breast carcinomas.

|  |  |  |  | Univariate 'ILC' HR+/HER2- |  |  | Univariate 'IDC' HR+/HER2- |  |  |
| --- | --- | --- | --- | --- | --- | --- | --- | --- | --- |
|  |  |  |  | N | HR [95%CI] | p-value | N | HR [95%CI] | p-value |
| DFS | TILs, Lymphoid score, median | high vs low |  | 360 | 0.82 [0.48-1.39] | 0.452 | 1875 | 0.66 [0.55-0.80] | **3.13E-05** |
| OS | TILs, Lymphoid score, median | high vs low |  | 361 | 1.19 [0.65-2.20] | 0.571 | 1876 | 0.71 [0.58-0.87] | **1.08E-03** |
| DFS, disease-free survival; OS, overall survival; N, effective; HR, hazards ratio; CI, confidence interval; bold is for significant p-value |  |  |  |  |  |  |  |  |  |

Altogether, these results suggest that the quality, more than the quantity, of immune cell infiltration should be studied to understand the differential immune contexture of HR^+^/HER2^-^ ILCs *vs* HR^+^/HER2^-^ IDCs.

### The low immune infiltrate differs qualitatively between HR^+^/HER2^-^ ILCs and HR^+^/HER2^-^ IDCs

We thus analyzed the composition and functional orientation of immune infiltrates in detail. First, we used Bindea’s metagenes to compare the enrichment of immune cell subsets between HR^+^/HER2^-^ ILCs and HR^+^/HER2^-^ IDCs ^27^. We focused on the most robust enrichments (P<1E-10) and found that the top three sub-populations enriched in HR^+^/HER2^-^ ILCs were CD8^+^ cytotoxic T cells (P=6.34E-21), eosinophils (P=1.85E-13), and mast cells (P=7.25E-32). Other subpopulations were also enriched in HR^+^/HER2^-^ ILCs, such as immature dendritic cells (P=1.66E-11) and dendritic cells (P=2.22E-18), cytotoxic cells (P=4.27E-17), B cells (P=2.65E-11), and T cells (P=8.40E-11), while HR^+^/HER2^-^ IDCs were strongly enriched in type 2 helper T cells (T_H_2, P=4.11E-21) (Figure 1C).

Second, and to confirm the enrichment more resolutely, we analyzed a publicly available scRNAseq dataset from HR^+^/HER2^-^ ILC and HR^+^/HER2^-^ IDC BCs ^18^, sorted for immune cells. We collected 13,447 cells from five patients: two with HR^+^/HER2^-^ IDCs, and three with HR^+^/HER2^-^ ILCs. We clustered the cells into 12 clusters, which were annotated by automatic label transfer annotation (Bindea, scAnnot and SingleR) (Supplementary Figure 2A-C) then refined manually based on the presence of canonical markers found in DEG (Figure 1D-E, Supplementary Table 6). Among the 12 clusters, only cluster_c8 consisted of residual contaminating epithelial cells, and expressed DEG such as *KRT19, KRT18, KRT8, EPCAM, CEACAM6,* and *CDH1.* As expected from the sample preparation, all the other clusters were of the leukocyte lineage. Six clusters were of the lymphocytic lineage. Cluster_c6, was a cluster of Natural Killer (NK) cells, characterized by DEG coding for NK canonical markers (*KLRF1*/*NKp80, NCR3/NKp30, NCR1*/*NKp46, FCGR3A*/*CD16*, *EOMES*), as well as granzymes (*GZMB, GZMA, GZMH, GZMM*), and granulysin (*GNLY*). Cluster_c0 was a cluster of CD8^+^ cytotoxic T cells, expressing canonical markers for CD8^+^ cytotoxic T cells (*CD8A, CD8B, CD3D, CD3E, CD3G, CD247, LCK, PTPRC, THEMIS, STAT4*), but also DEG encoding for granzymes (*GZMK, GZMH, GZMA, GZMM, GZMB*), perforin (*PRF1*) and granulysin (*GNLY*). Cluster_c3 was a cluster of helper-like progenitor CD8^+^ cytotoxic T cells, close to cluster_c0 (*GZMK, CD8A, CD8B etc.*), but with higher expression of *ICOS*, *CD40L*, *IL2R* and *IL2*. They are not classical CD8^+^ cytotoxic killer cells, and rather function as immunoregulatory cells, providing support and being a reservoir of functional CD8^+^ cytotoxic T cells. Cluster_c1 was a cluster of activated/follicular helper T/helper T cells, characterized by DEG that are canonical markers of T cells (*CD3E, CD3G, CD3D, CD28, ICOS, CD40L*). It also expressed the *GATA3* transcription factor, suggesting a polarization in T_H_2 cells. Cluster_c2 was a cluster of naive/memory T cells, characterized by DEG such as *SELL, CCR7, CD3E, CD40LG*. Cluster_c5 was a cluster of B cells, characterized by DEG that are canonical markers of B cells: *CD79A, CD79B, IGHM, IGKC, IGHD*, coding for Immunoglobulin alpha, *MS4A1/CD20*, *CD19*, *CD22*, *POU2F2*, and *PAX5*.

Five clusters were of the myeloid lineage. Cluster_c4 was a cluster of type 2 tumor-associated macrophages (TAM2) characterized by DEG involved in complement cascade (*C1QC, C1QA, C1QB, C3*), and antigen presentation (*CD74*/*CLIP*, *CIITA*, *HLA-DRB1, HLA-DPA1, HLA-DPB1, HLA-DQB1, HLA-DQA1, HLA-DMA, HLA-DMB, HLA-DQA2, HLA-DOA, HLA-DQB2*). It also expressed canonical markers of macrophages such as *CD68*, *CD14*, *MSR1/CD204*, *MRC1/CD206,* but also *CD163* suggesting a TAM2 polarization. Cluster_c7 was quite similar to cluster_c4. It was another TAM2 subtype expressing DEG involved in complement cascade (*CFD, CFP, CR1, C5AR1*) and antigen presentation (*HLA-DRB5, HLA-DRB1, HLA-DMA, HLA-DMB, CIITA*), but also canonical markers of macrophages such as *CD68, CD14, ITGAX/CD11c, ITGAM/CD11b, TLR4, MARCO,* but also *CD163* suggesting a TAM2 polarization. Both clusters expressed *IL1B, HIF1A, ENTPD1* and *PTGS2* but cluster_c4 was associated with expression of *APOE* and *SPP1* suggesting a lipid-associated TAM2, whereas cluster_7 expressed *TGFB1* suggesting an immunosuppressive TAM2. Similarly to clusters_c7 and _c4, the cluster_c9 expressed canonical markers of macrophages (*CD68*, *ITGAX*/*CD11c*, *HLA*), but also expressed *POU2F2*, suggesting a state close to monocyte (annotated monocyte-like macrophages). Cluster_c10 was a cluster of dendritic cells, characterized by DEG such as *CD1C, CD1E, CD1B, CD1D, CD86, MRC1, ITGAX*/*CD11c*, *CD68, IRF8* and *IRF4*, or involved in antigen presentation (*HLA-DQA1, HLA-DQA2, HLA-DRB5, HLA-DQB1, HLA-DMB, HLA-DMA, HLA-DPB1, HLA-DRA, HLA-DRB1, HLA-DPA1, HLA-DQB2, HLA-DOA, HLA-DOB, CIITA, CD74/CLIP*). Cluster_c11 was a cluster of mast cells, characterized by expression of tryptases (*TPSAB1, TPSB2, TPSD1*) but also *ITGAM, CD33, KIT, FCER1A*, that are canonical markers.

To confirm at the single-cell level the enrichment of immune cells observed in bulk data, we compared clusters enrichment between HR^+^/HER2^-^ ILCs and HR^+^/HER2^-^ IDCs. Due to the small number of cases in the single-cell dataset, no significant enrichment could be established. Nevertheless, HR^+^/HER2^-^ ILCs tended to be enriched in cells corresponding to the cluster_c0 of CD8^+^ cytotoxic T cells (23.1% *vs* 11.5%, respectively), the cluster_c1 of helper T cells (19.9% *vs* 8.8%), and the cluster_c2 of naive/memory T cells (18.5% *vs* 7.4%), but not in cluster_c3 of helper-like progenitor CD8^+^ cytotoxic T cells which are immune organizers and a reservoir for CD8^+^ cytotoxic T cells (12.7% *vs* 14.8%). Moreover, HR^+^/HER2^-^ IDCs tended to be enriched in cluster_c4 of TAM2 macrophages (39.4% *vs* 3.0%, respectively) (Figure 1F-H). Altogether, the scRNAseq data confirmed the enrichment of CD8^+^ cytotoxic T cells in HR^+^/HER2^-^ ILCs, as observed in the bulk data. It also reinforced the enrichment of T cells in HR^+^/HER2^-^ ILCs. B cells cluster_c5 seemed to be slightly more present in HR^+^/HER2^-^ ILCs than in HR^+^/HER2^-^ IDCs, following the same tendency as the bulk results. We also examined the patient contributions to the TAM2 cluster_c4, which seems to yield a more unexpected result, but found that most macrophages originated from a single patient. Therefore, this finding could not be interpreted further with this dataset due to an obvious representativity bias (Supplementary Figure 2D). Finally, mast cells and dendritic cells were too few to assess an enrichment.

Altogether, both bulk data (providing strong statistical power) and single-cell data (providing cell subtype precision) tend to show that CD8^+^ cytotoxic T cells are present in HR^+^/HER2^-^ ILCs in higher proportion than in HR^+^/HER2^-^ IDCs.

### The anti-tumor immune infiltrate of HR^+^/HER2^-^ ILCs is better organized and is associated with a higher chance of response to immune checkpoint inhibitors

Because CD8^+^ cytotoxic T cells are key to mediate anti-tumor immune functions, we searched for signs of immunogenicity and lymphocyte priming. We first evaluated the signature for TLS ^29^ and found a small enrichment in HR^+^/HER2^-^ ILCs (P=3.55E-3) (Table 3). We next sought to confirm the presence of organized lymphoid structures by assessing CD3^+^ and CD20^+^ lymphocyte aggregates using IHC (34 HR^+^/HER2^-^ ILCs and 36 HR^+^/HER2^-^ IDCs) applied on tumor slide (one per patient). Two different types of B and T cell aggregates were found: “unorganized” or “organized”, the latter being known as TLS-like structure; and among the TLS-like structures, two subtypes of “immature” or “mature” TLS were defined based on the spatial organization and density of aggregates ^40^ (Figure 2A). TLS-like were enriched in HR^+^/HER2^-^ ILCs compared to HR^+^/HER2^-^ IDCs (P=7E-3, Figure 2B), but this enrichment was mainly due to immature TLS (P=7E-3), while the proportion of mature TLS-like structures was not different between HR^+^/HER2^-^ ILCs and HR^+^/HER2^-^ IDCs (P=0.2, Figure 2C). While the presence of a single TLS may be attributed to “chance”, the occurrence of multiple TLS indicates a specific immune mobilization. Thus, we searched for tumors sections presenting more than two TLS-like. We found that they were more frequently observed in HR^+^/HER2^-^ ILCs (P=1E-3) (Figure 2D). The higher presence of TLS is also supported by the enrichment in follicular helper T cells in HR^+^/HER2^-^ ILCs (P=1.29E-8, Figure 1C).

**Figure 2:**
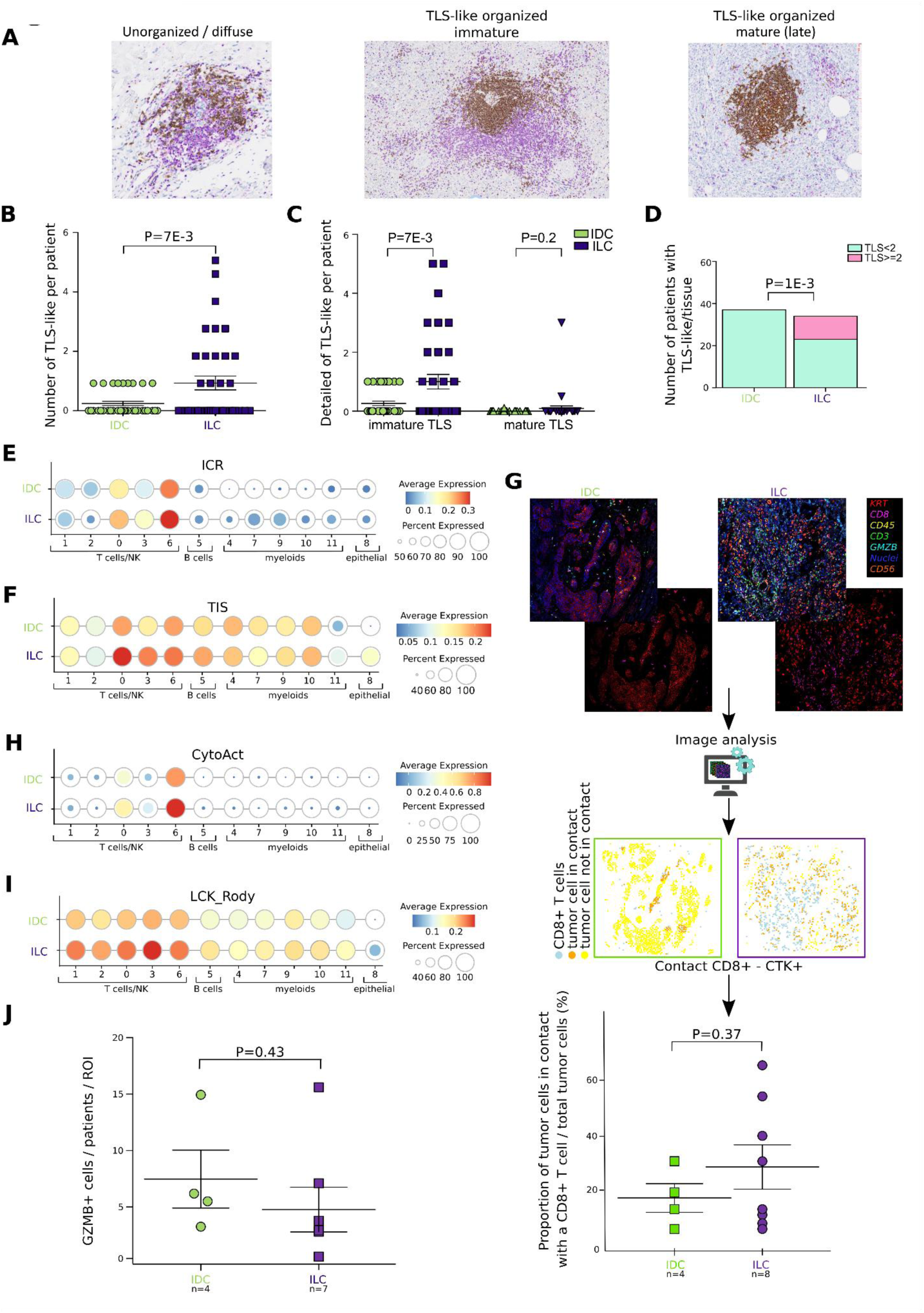
HR^+^/HER2^-^ ILCs demonstrated a mobilized and organized anti-tumor potential compared to HR^+^/HER2^-^ IDCs. (A) Immunohistochemistry pictures of the different classification of B cells (CD20^+^, brown) and T cells (CD3^+^, purple) aggregates; “unorganized/diffuse” or “TLS-like organized mix immature/ma-ture(late)”. TLS; tertiary lymphoid structures. (B) Quantification of the number of TLS-like *per* patients in n=34 HR^+^/HER2^-^ ILCs *vs* n=36 HR^+^/HER2^-^IDCs. P-value was obtained for Student t-test. Green is for IDCs and purple is for ILCs and this color code will be used throughout the article. (C) Detailed quantification of TLS-like *per* patients, between immature and mature subtypes, in n=34 HR^+^/HER2^-^ ILCs *vs* n=36 HR^+^/HER2^-^ IDCs. P-value was obtained for Student t-test. (D) Quantification of the number of patients with two or more than two TLS-like (pink) or less than two TLS-like (blue) *per* tissue in n=34 HR^+^/HER2^-^ ILCs *vs* n=36 HR^+^/HER2^-^ IDCs. P-value was obtained for contingency Fisher’s test. (E) Dot plots representing the enrichment of the ICR signature in clusters of the scRNAseq GSE193911 dataset (Onkar *et al*) in HR^+^/HER2^-^ ILCs *vs* HR^+^/HER2^-^ IDCs. The size of the circle represents the per-cent of cells expressing the signature and the color represent the absolute strength of the expres-sion. (F) Dot plots representing the enrichment of the TIS signature in clusters of the scRNAseq GSE193911 dataset (Onkar *et al*) in HR^+^/HER2^-^ ILCs *vs* HR^+^/HER2^-^ IDCs. The size of the circle represents the per-cent of cells expressing the signature and the color represent the absolute strength of the expres-sion. (G) Workflow & quantification of the proportions of tumor cells in contact with a cytotoxic T cell. Mul-tiplexed immunofluorescence (up panel) identified several immune subtypes (tumor; KRT^+^, cyto-toxic T cell; CD8^+^, immune cells; CD45^+^, T cell; CD3^+^; active cytotoxic cells; GZMB^+^, Natural killer; CD56^+^) and cellular neighborhood analysis (mid panel) quantified tumor cells in contact (*orange*) or not (*yellow*) with a CD8^+^ T cell (*light blue*) differentially in 12 patients (n=8 HR^+^/HER2^-^ ILCs and n=4 HR^+^/HER2^-^ IDCs). Created in BioRender. Mamessier, E. (2026) https://BioRender.com/ x3x0w33. Box plot (down panel) represents the proportion of tumor cells in contact with a CD8^+^ T cell in HR^+^/HER2^-^ ILCs *vs* HR^+^/HER2^-^ IDCs. P-value was obtained for Student t-test. n; effective. (H) Dot plots representing the enrichment of the CytoAct signature in clusters of the scRNAseq GSE193911 dataset (Onkar *et al*) in HR^+^/HER2^-^ ILCs *vs* HR^+^/HER2^-^ IDCs. The size of the circle repre-sents the percent of cells expressing the signature and the color represent the absolute strength of the expression. (I) Dot plots representing the enrichment of the LCK signature in clusters of the scRNAseq GSE193911 dataset (Onkar *et al*) in HR^+^/HER2^-^ ILCs *vs* HR^+^/HER2^-^ IDCs. The size of the circle represented the percent of cells expressing the signature and the color represent the absolute strength of the ex-pression. (J) Quantification the number of GZMB+ cells *per* patient *per* region of interest (ROI) normalized by, differentially between n=7 HR^+^/HER2^-^ ILCs and n=4 HR^+^/HER2^-^ IDCs. P-value is for Student t-test. n; effective.

**Table 3:** Enrichment in functional anti-tumor or immunosuppressive molecular profile in HR+/HER2- invasive lobular or ductal breast carcinomas.

| Signature | N | ILC | IDC | p-value | Status |
| --- | --- | --- | --- | --- | --- |
| TLS_GES_Coppola_2011 | 3046 | -0.06 (-2.01-1.5) | -0.15 (-2.57-2.44) | **3.55E-03** | Up ILC |
| ICR_score_Bertucci_2018 | 3046 | -0.01 (-1.87-1.96) | -0.12 (-2.31-3.13) | **1.69E-03** | Up ILC |
| TIS_GES_Ayers_2017 | 3046 | 0 (-2.05-2.07) | -0.14 (-2.46-2.78) | **5.67E-05** | Up ILC |
| CytoAct_Rooney_2015 | 3046 | 0.01 (-2.13-3.85) | -0.09 (-3.02-4.65) | **3.61E-02** | Up ILC |
| LCK_GES_Rody_2009 |  |  |  | **4.42E-04** | up IDC |
| Low | 2413 | 325 (73%) | 2088 (80%) |  |  |
| High | 633 | 121 (27%) | 512 (20%) |  |  |
| IFNalpha_Gatza_2011 | 3046 | 0.42 (0.01-0.99) | 0.47 (0.01-0.99) | **1.41E-03** | up IDC |
| IFNgamma_Gatza_2011 | 3046 | 0.41 (0.01-0.98) | 0.46 (0-0.99) | **1.94E-04** | up IDC |
| APMS_score | 3046 | -3.09 (-25.38-25.9) | -0.08 (-37.84-31.62) | **1.19E-13** | up IDC |
| Hypoxia_Gatza_2011 | 3046 | 0.55 (0.02-0.97) | 0.48 (0-1) | **7.50E-08** | Up ILC |
| Acidosis_Gatza_2011 | 3046 | 0.63 (0.01-0.98) | 0.48 (0.01-0.95) | **1.10E-47** | Up ILC |
| TGFB3 | 3046 | 0.85 (-2.27-2.68) | 0.47 (-2.53-2.88) | **4.24E-23** | Up ILC |
| TGFB2 | 3046 | 0.02 (-3.32-5.25) | -0.25 (-4.15-14.6) | **2.85E-05** | Up ILC |
| TGFB1 | 3045 | 0.57 (-2.26-3.8) | 0.43 (-3.26-3.61) | **2.37E-03** | Up ILC |
| PTGDS | 4476 | 8.68 (5.29-13.86) | 9.21 (6.66-13.82) | **1.68E-24** | Up ILC |
| PTGS2 | 4477 | 3.79 (0.55-11.19) | 4.09 (0.95-9.84) | **2.54E-09** | Up ILC |
| PTGER4 | 4477 | 7.12 (2.1-11.54) | 7.64 (3.23-10.37) | **3.49E-26** | Up ILC |
| *N, effective; GES, gene expression signature; bold is for significant p-value* |  |  |  |  |  |

We then tested two other signatures predictive of favorable response to ICIs: the ICR and the TIS signatures ^19^ ^41^. Both were stronger in HR^+^/HER2^-^ ILCs than in HR^+^/HER2^-^ IDCs (P=1.69E-3, and P=5.67E-5, respectively, Table 3). In the scRNAseq dataset, we confirmed the enrichment of these two signatures in the dysfunctional CD8^+^ cytotoxic T cell cluster_c0, helper-like progenitor CD8^+^ cytotoxic T cells cluster_c3, and NK cluster_c6, of HR^+^/HER2^-^ ILC tumors, compared to HR^+^/HER2^-^ IDC tumors (Figure 2E-F).

We also investigated the physical accessibility of tumor cells to cytotoxic T cells by examining the spatial distribution of tumor cells (pan-KRT^+^) compared to cytotoxic T cells (CD8^+^), using multiplex-IF in 12 patients (6 per group). The proportion of tumor cells in close proximity (less than 25µm) to a CD8^+^ cytotoxic T cell tend to be higher in HR^+^/HER2^-^ ILCs (∼50%) than in HR^+^/HER2^-^ IDCs (<25%) (Figure 2G), thus supporting better physical accessibility of CD8^+^ cytotoxic T cells to tumor cells in HR^+^/HER2^-^ ILCs.

Altogether, these results suggest that the anti-tumor immune infiltrate in HR^+^/HER2^-^ ILC tumors can interact with tumors cells, is more organized and may have a higher potential to respond to ICIs than HR^+^/HER2^-^ IDC tumors. Therefore, we examined the functional activation of immune infiltrates.

### Anti-tumor response in HR^+^/HER2^-^ ILCs is poorly effective

To further explore the functionality of immune infiltrates, we assessed the enrichment for several immune activation signatures. First, we examined the CytoAct signature, a score for cytotoxic activation ^31^. This score was higher in HR^+^/HER2^-^ ILCs than in HR^+^/HER2^-^ IDCs (P=3.61E-2) (Table 3). At the single-cell level, this enrichment in HR^+^/HER2^-^ ILCs was mainly observed in cluster_c6 of NK cells, and to a lesser extent in cluster_c0 of CD8^+^ cytotoxic T cells (Figure 2H). The fact that CytoAct was more enriched in NK cluster_c6 compared to CD8^+^ cytotoxic T cells cluster_c0 suggested a dysfunctionality of CD8 cytotoxic cells in both HR^+^/HER2^-^ ILCs and HR^+^/HER2^-^ IDCs. This is also evidenced with the expression of cytotoxic genes shown in Figure 1E.

We also found that HR^+^/HER2^-^ ILCs were enriched with the LCK kinases signature comparatively to HR^+^/HER2^-^ IDCs (P=4.42E-4, Table 3); this is a signature of T-cell receptor (TCR) activation signaling that further promotes proliferation while restraining effector differentiation ^42^ ^32^. We confirmed enrichment of this signature in HR^+^/HER2^-^ ILCs within lymphocytes, most strongly in cluster_c3 of helper-like progenitor CD8^+^ cytotoxic T cells and to a lesser extent in cluster_c0 CD8^+^ cytotoxic T cells and cluster_c6 NK cells, which are the clusters of effector cells (Figure 2I).

We then evaluated the expression of cytokines characteristic of an efficient anti-tumor response. Notably, we found that the Gatza’ signatures of IFNα and IFNγ pathways activation ^43^ were less enriched in HR^+^/HER2^-^ILCs than in HR^+^/HER2^-^ IDCs (P=1.41E-3 and P=1.94E-4 respectively, Table 3).

Finally, we examined the effective antitumor immune response, represented by granzyme degranulation, using mtpx IF, and found no significant enrichment in favor of HR^+^/HER2^-^ ILCs but rather a tendency toward HR^+^/HER2^-^ IDCs (P=0.8, Figure 2J).

Altogether, our results show that, despite a higher pro-cytotoxic activation potential, the CD8^+^ cytotoxic T cells in HR^+^/HER2^-^ ILCs show signs of dysfunction or improper activation. Our observation suggests that a mechanism blocks the final activation of anti-tumor immune response in HR^+^/HER2^-^ ILCs.

### CD8^+^ cytotoxic T cells have not transitioned to a fully activated stage in HR^+^/HER2^-^ ILCs

To understand the extent of this defect in T cell activation in HR^+^/HER2^-^ ILCs compared to HR^+^/HER2^-^ IDCs, we examined lymphocyte maturation status using CIBERSORT, a versatile computational method for quantifying cell fractions from bulk tissue gene expression profiles. This approach provides another granularity regarding lymphocyte maturation stages compared to the previously used Bindea’s method. We found that HR^+^/HER2^-^ ILCs were enriched in CD8^+^ T cells (P=1.08E-11), consistent with previous results, but also with memory B cells (P=3.16E-7) and CD4^+^ memory resting T cells (P=1.86E-3). However, CD4^+^ memory T cells with an activated status, and regulatory T cells were enriched in HR^+^/HER2^-^ IDCs (P=1.84E-3 and P=4.28E-6, respectively) (Figure 3A).

**Figure 3:**
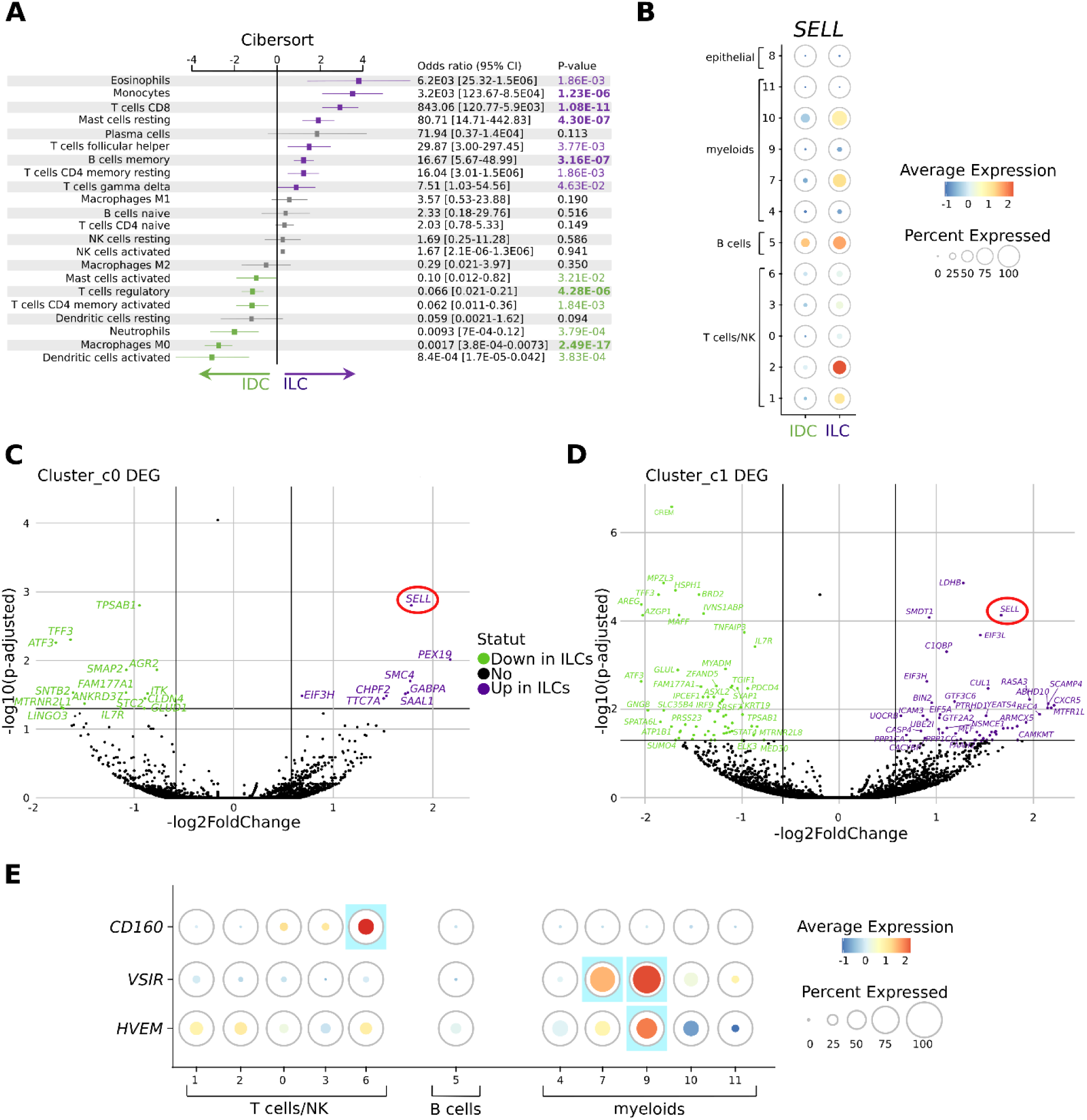
HR^+^/HER2^-^ ILCs tumors are deprived of infiltrating effector T cells and effective antigen presenting cells. (A) Tree plot representing the enrichment of CIBERSORT immune score in HR^+^/HER2^-^ ILCs *vs* HR^+^/HER2^-^IDCs from bulk RNA sequencing datasets. CI; confidence interval, M0, M1 and M2; type 0, 1 2 mac-rophages subtypes. P-value in purple referred to p-value enriched in HR^+^/HER2^-^ ILCs, and green ones for p-value enriched in HR^+^/HER2^-^ IDCs. Grey frames are for readability. Bold is for P-value < 1E-5. Green is for IDCs and purple is for ILCs and this color code will be used throughout the article. (B) Dot plots representing the enrichment of *SELL* in clusters of the scRNAseq GSE193911 dataset (On-kar *et al*) in HR^+^/HER2^-^ ILCs *vs* HR^+^/HER2^-^ IDCs. The size of the circle represents the percent of cells expressing the signature and the color represent the relative strength of the expression. (C) Volcano plot representing differentially expressed genes (DEG) of the cluster_c0 from scRNAseq datasets in HR^+^/HER2^-^ ILCs *vs* HR^+^/HER2^-^ IDCs. Green-colored dots represented genes significantly enriched in HR^+^/HER2^-^ IDCs whereas purple-colored ones represented genes significantly enriched in HR^+^/HER2^-^ ILCs. *SELL* was highlighted by a red circle. (D) Volcano plot representing differentially expressed genes (DEG) of the cluster_c1 from scRNAseq datasets in HR^+^/HER2^-^ ILCs *vs* HR^+^/HER2^-^ IDCs conditions. Green-colored dots represented genes significantly enriched in HR^+^/HER2^-^ IDCs whereas purple-colored ones represented genes signifi-cantly enriched in HR^+^/HER2^-^ ILCs. *SELL* was highlighted by a red circle. (E) Dot plots representing the enrichment of *CD160*, *VSIR*, *HVEM* in clusters of the scRNAseq GSE193911 dataset (Onkar *et al*) in HR^+^/HER2^-^ ILCs *vs* HR^+^/HER2^-^ IDCs. The size of the circle repre-sents the percent of cells expressing the signature and the color represent the relative strength of the expression. Blue frame highlighted the clusters enriched in genes of interest.

In parallel, we confirmed the non-effector status of T cells and B cells infiltrating HR^+^/HER2^-^ ILCs in the scRNAseq dataset by showing that naive/memory T cells cluster_c2 and B cells cluster_c5 overexpressed *SELL*, a molecule characteristic of naive and memory lymphocytes that have not reached the effector stage (Figure 3B). Interestingly, when we examined the DEG for each cluster between HR^+^/HER2^-^ ILCs and HR^+^/HER2^-^ IDCs, we found that *SELL* was the top DEG in cluster_c0 (CD8^+^ cytotoxic T cells), and the sixth top DEG in cluster_c1 (activated / follicular helper T cells / helper T cells) (P=1.58E-3 and P=7.4E-5 respectively, Figure 3C-D and Supplementary Table 7) ; in each cluster, it was overexpressed in HR^+^/HER2^-^ ILCs *vs* HR^+^/HER2^-^ IDCs, again consistent with a non-effector status for these cells in HR^+^/HER2^-^ ILCs. Altogether, our results show that T cells had not transitioned to a full effector phenotype in HR^+^/HER2^-^ ILCs.

Secondly, for the non-naive T cells, we specifically analyzed their potential level of exhaustion or inhibition by examining the expression of several immune checkpoints (inhibitors or activators) and their ligands (Table 4). We found that HR^+^/HER2^-^ ILCs were enriched in inhibitory molecules expressed by T cells, such as *PDCD1*/*PD-1* (P=2.20E-02) and *PVRIG* (P=9.65E-15) (Table 4) ^44^ ^45^. However, we did not find enrichment for other conventional molecules expressed in exhausted T cells (*LAG3*, *HAVCR2*). This suggests that the overall activation of lymphocytes infiltrating ILC tumors results from a dysfunctional profile or improper activation, rather than an exhausted phenotype.

**Table 4:**
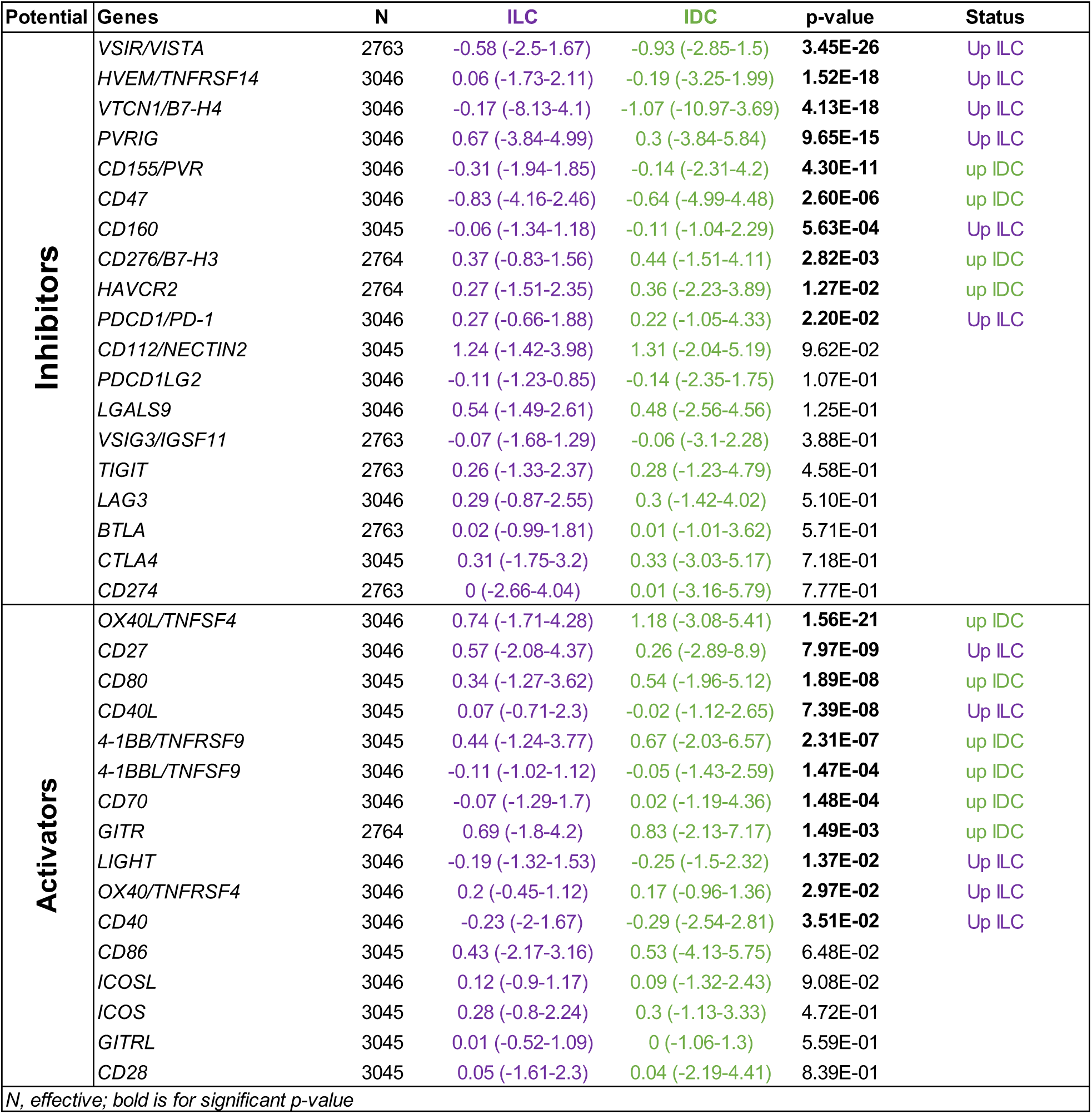
Expression of immune checkpoint receptors and ligands in HR+/HER2- invasive lobular or ductal breast carcinomas.

In line with this, HR^+^/HER2^-^ ILCs also expressed inhibitory molecules that can act in *cis* or in *trans* on T-cell activation, such as *CD160* (P=5.63E-4), mainly expressed by NK cells ^46^, as well as *VSIR*/*VISTA* (P=3.45E-26), mainly expressed by myeloid cells, but also found on T cells ^47^, and *HVEM*/*TNFRSF14* (P=1.52E-18), which is expressed by tumor cells and resting leukocytes ^48^ ^49^. In addition, HR^+^/HER2^-^ ILCs were enriched in inhibitory molecules, notably *VTCN1*/*B7-H4* (P=4.13E-18), which is mainly expressed by the tumor cells, impairing cytotoxic T cell activity ^50^ and representing a new target for immunotherapy ^51^ (Table 4).

We looked at the expression of molecules acting in *cis*- or in *trans*- to assess their expression across different immune cell compartments in the single-cell dataset. We found that *CD160* was mainly expressed in cluster_c6 of NK cells, consistent with the literature. We detected a strong expression of *VSIR* in the myeloid compartment, notably in cluster_c7 TAM2 and cluster_c9 monocyte-like macrophages, in agreement with previous reports. In addition to the slight expression of *HVEM* in T cells (cluster_c1 and cluster_c0), NK cells (cluster_c6), and TAM2 (cluster_c7), we found that *HVEM* was strongly expressed by monocyte-like macrophages (cluster_c9), as expected ^52^ (Figure 3E). The high expression of *VSIR* and *HVEM* in the myeloid compartment are potential candidates for T cells inhibition by macrophages. Furthermore, we paid a specific attention to *VTCN1,* a potent inhibitor of T cell functions (activation, proliferation, cytokine production…). VTCN1 can be expressed by either epithelial tumor cells or antigen-presenting cells. Because the single-cell dataset of Onkar *et al* lack of epithelial cells, we collected a second single-cell dataset for which both myeloid and tumor cells were available ^35^. The analysis of 1 ILC *vs* 10 IDCs revealed that luminal epithelial tumor cells were the primary source of VTCN1/B7-H4 compared to myeloid cells (Supplementary Figure 2E). To complete, we turned back to our analysis of tumor bulk samples. *VTCN1* was among the top DEG in HR^+^/HER2^-^ ILCs compared to HR^+^/HER2^-^ IDCs (Supplementary Figure 1B, Supplementary Table 4).

Interestingly, few inhibitory immune checkpoint molecules were overexpressed in HR^+^/HER2^-^ IDCs, such as *CD47* (P=2.6E-6) and *CD155/PVR* (P=4.3E-11) and to a lower extent *HAVCR2* (P=1.27E-2) and *CD276/B7-H3* (P=2.82E-3) (Table 4). Inversely, several immune activator molecules were found to be increased in HR^+^/HER2^-^ IDCs, including *GITR* (P=1.49E-3), *OX40L* (P=1.56E-21), *4-1BB* (P=2.31E-7), *4-1BBL* (P=1.47E-4), *CD80* (P=1.89E-8), and *CD70* (P=1.48E-4), while *CD40* (P=3.51E-2), *CD40L* (P=7.39E-8), *LIGHT* (P=1.37E-2), *OX40/TNFRSF4* (P=2.97E-2) and *CD27* (7.97E-9) were enriched in HR^+^/HER2^-^ ILCs but globally with weaker p-values (Table 4).

Altogether, the more frequent and significant overexpression of ICIs and the less frequent and significant overexpression of immune checkpoint activators in HR^+^/HER2^-^ ILCs *vs* HR^+^/HER2^-^ IDCs suggest an “unproper” yet still immunosuppressed effector profile of T cells in in HR^+^/HER2^-^ ILCs.

### HR^+^/HER2^-^ ILCs evidenced a defective antigen-presenting compartment

Because CD8^+^ cytotoxic T lymphocytes in the tumor microenvironment can be reactivated by local antigen-presenting cells (such as macrophages or dendritic cells), we looked at the myeloid compartment, noting that HR^+^/HER2^-^ ILCs were less infiltrated by myeloid cells than HR^+^/HER2^-^ IDCs (Figure 1B). Regarding professional antigen-presenting cells (APC), type 0 macrophages were highly enriched in HR^+^/HER2^-^ IDCs compared to HR^+^/HER2^-^ ILCs (P=2.49E-17) (Figure 3A). These cells are macrophage precursors in a resting state, before polarization into type 1 or type 2 macrophages. This lack of polarization is consistent with the overexpression of *CD47* (P=2.6E-6), a key inhibitor of myeloid cell activation (Table 4). Except for this progenitor subtype, the macrophage compartment did not show major differences between HR^+^/HER2^-^ ILCs and HR^+^/HER2^-^ IDCs.

Immature dendritic cells were enriched in HR^+^/HER2^-^ ILCs (P=1.66E-11) (Figure 1C), while HR^+^/HER2^-^ IDCs were enriched in dendritic cells with an activated status (P=3.83E-4) (Figure 3A). This observation is reinforced by a stronger expression of genes coding for the antigen-presentation machinery, measured by the AP-MS score, in HR^+^/HER2^-^ IDCs (P=1.19E-13, Table 3).

Altogether, these data suggest that mature antigen presentation ability, required for proper activation of anti-tumor immunity, is defective in most APCs from HR^+^/HER2^-^ ILCs, and might contribute to the incompletely activated phenotype of CD8^+^ cytotoxic T cells.

### HR^+^/HER2^-^ ILCs have a stronger immunosuppressive environment

In parallel, we also investigated other potential sources of immune deficiency that could contribute to this lack of activation, such as hypoxia, acidosis, inhibitory cytokines or inflammatory molecules. First, we found a strong enrichment of Gatza’s hypoxia and particularly acidosis signatures ^43^ in HR^+^/HER2^-^ ILCs compared to HR^+^/HER2^-^ IDCs (P=7.5E-8 and P=1.1E-47 respectively) (Table 3). Acidosis causes a low pH in the tumor, which further degrades the matrix, facilitating tumor cell migration, inhibits cytotoxic effector cells and prevents APC activity. Second, we analyzed the expression of immunosuppressive and pro-tumorigenic TGFβ family of cytokines and observed strong enrichment of *TGFB3*, *TGFB2*, and *TGFB1* in HR^+^/HER2^-^ ILCs (P=4.24E-23, P=2.85E-5 and P=2.37E-3 respectively). Additionally, we also looked at genes involved in prostaglandin synthesis, previously described as pro-inflammatory and immunoregulatory molecules. *PTGDS*, *PTGS2* and *PTGER4* were enriched in HR^+^/HER2^-^ ILCs (P=1.68E-24, P=2.54E-9 and P=3.49E-26, respectively) (Table 3). Altogether, these results suggest that the microenvironment of HR^+^/HER2^-^ ILCs is acidic, pro-inflammatory and pro-tumoral.

### HR^+^/HER2^-^ ILCs are enriched in inflammatory cancer-associated fibroblasts and pericyte-like subsets compared to HR^+^/HER2^-^ IDCs

The source of pro-tumoral inflammation can have multiple origins, but since the main enrichment between HR^+^/HER2^-^ ILCs and HR^+^/HER2^-^ IDCs involved stromal cells (Figure 1B), we focused on these cells. We aimed to precise the quality of stromal population enrichment by investigating enrichment in neuron, fibroblast, endothelial cell, and adipocyte signatures. Except for endothelial cells, most signatures were indeed enriched in HR^+^/HER2^-^ ILCs (Figure 4A), and the potential involvement of these cell types deserves further attention. Because fibroblasts are quantitatively the most prominent component of the stroma ^53^ ^54^, and because multiple subtypes exist with various functions that can affect anti-tumor response, we investigated the quality of this compartment ^55^ ^56^. We evaluated the enrichment of known cancer-associated fibroblast (CAF) signatures in HR^+^/HER2^-^ ILCs and HR^+^/HER2^-^ IDCs ^57^ ^55^ ^56^. We observed that HR^+^/HER2^-^ ILCs were significantly enriched in immunoregulatory CAFs (iCAF; P=1.4E-37), and specifically in the IL-iCAF subtype (interleukin-signaling pathway CAF; P=1.22E-49) and the detox-iCAF subtype (detoxification pathway CAF; P=8.85E-48) (Figure 4B). Detox-iCAF and IL-iCAF are both described as immunoregulatory CAFs involved in lymphocyte recruitment, through the production of IL6, CXCL12 and other inflammatory factors ^60^. Individually, the differential enrichment in HR^+^/HER2^-^ ILCs *vs* HR^+^/HER2^-^ IDCs was stronger for iCAF signature compared to myofibroblastic CAF (myCAF) signature (P=3.54E-35 *vs* P=3.86E-3) and the iCAF/myCAF signature ratio was higher in HR^+^/HER2^-^ ILCs (P=3.73E-30) (Figure 4C-D). While iCAFs typically promote an immunosuppressive microenvironment by secreting cytokines such as IL-6 and CXCL12, which recruit regulatory T cells and inhibit T-cell activity, myCAF contributes to build physical barriers *via* dense extracellular matrix deposition to limits immune cell infiltration and cytotoxicity ^61^ ^62^ ^63^. Consistent with iCAFs enrichment, *IL6*, *CXCL12*, *IL33*, *CXCL2*, *CFD*, *LIF* and *TGFB3* were among the most frequent soluble factors correlating with CAF presence, ranking in the top 10 most highly expressed soluble factors in HR^+^/HER2^-^ ILCs (Supplementary Table 8). Additionally, enrichment in IL-iCAF metagene strongly correlated with genes characteristic of CAFs, including the inflammatory genes *ANXA1* (Pearson r=0.79, P=1.37E-94), *CXCL12* (Pearson r=0.79, P=1.57E-94), *IL33* (Pearson r=0.76, P=3.3E-85), *IL6* (Pearson r=0.64, P=4.33E-52), and *PDGFRA* (Pearson r=0.72, P=4.14E-71) (Figure 4E-F, Supplementary Table 8).

**Figure 4:**
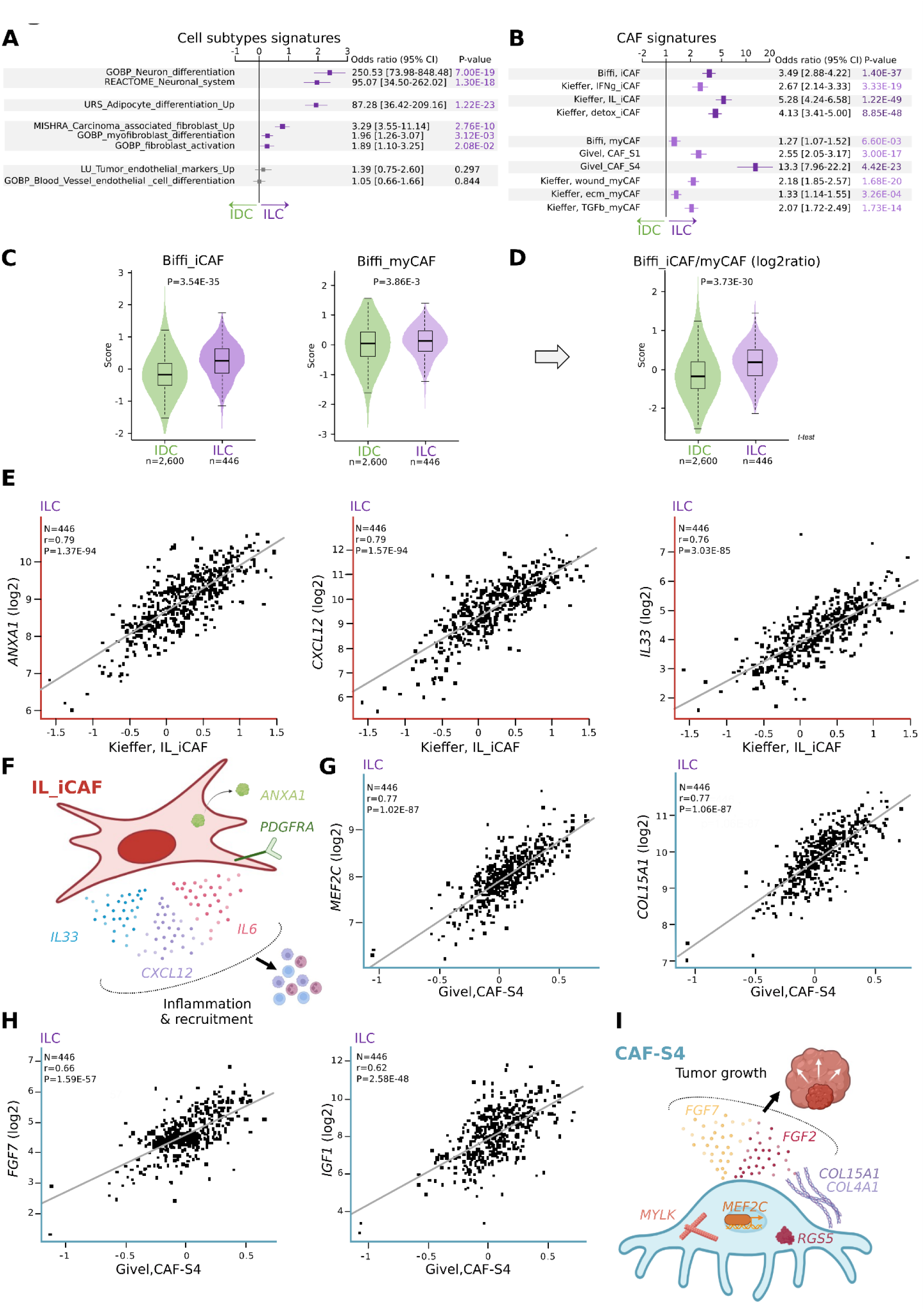
HR^+^/HER2^-^ ILCs tumors are deprived of infiltrating effector T cells and effective antigen presenting cells. (A) Tree plot representing the enrichment of CIBERSORT immune score in HR^+^/HER2^-^ ILCs *vs* HR^+^/HER2^-^IDCs from bulk RNA sequencing datasets. CI; confidence interval, M0, M1 and M2; type 0, 1 2 mac-rophages subtypes. P-value in purple referred to p-value enriched in HR^+^/HER2^-^ ILCs, and green ones for p-value enriched in HR^+^/HER2^-^ IDCs. Grey frames are for readability. Bold is for P-value < 1E-5. Green is for IDCs and purple is for ILCs and this color code will be used throughout the article. (B) Dot plots representing the enrichment of *SELL* in clusters of the scRNAseq GSE193911 dataset (On-kar *et al*) in HR^+^/HER2^-^ ILCs *vs* HR^+^/HER2^-^ IDCs. The size of the circle represents the percent of cells expressing the signature and the color represent the relative strength of the expression. (C) Volcano plot representing differentially expressed genes (DEG) of the cluster_c0 from scRNAseq datasets in HR^+^/HER2^-^ ILCs *vs* HR^+^/HER2^-^ IDCs. Green-colored dots represented genes significantly enriched in HR^+^/HER2^-^ IDCs whereas purple-colored ones represented genes significantly enriched in HR^+^/HER2^-^ ILCs. *SELL* was highlighted by a red circle. (D) Volcano plot representing differentially expressed genes (DEG) of the cluster_c1 from scRNAseq datasets in HR^+^/HER2^-^ ILCs *vs* HR^+^/HER2^-^ IDCs conditions. Green-colored dots represented genes significantly enriched in HR^+^/HER2^-^ IDCs whereas purple-colored ones represented genes signifi-cantly enriched in HR^+^/HER2^-^ ILCs. *SELL* was highlighted by a red circle. (E) Dot plots representing the enrichment of *CD160*, *VSIR*, *HVEM* in clusters of the scRNAseq GSE193911 dataset (Onkar *et al*) in HR^+^/HER2^-^ ILCs *vs* HR^+^/HER2^-^ IDCs. The size of the circle repre-sents the percent of cells expressing the signature and the color represent the relative strength of the expression. Blue frame highlighted the clusters enriched in genes of interest.

In addition to iCAF subsets, the pericyte-like subset (CAF-S4) was the most enriched subtype of myCAFs in HR^+^/HER2^-^ ILCs (P=4.42E-23) (Figure 4B). It strongly correlated with the myofibroblast genes *MEF2C* (Pearson r=0.77, P=1.02E-87), *COL15A1* (Pearson r=0.77, P=1.06E-87), *COL4A1* (Pearson r=0.55, P=6.18E-36), *MYLK* (Pearson r=0.58, P=8.88E-42), but also, as expected, the pericyte marker *RGS5* (Pearson r=0.50, P=6.75E-30) (Figure 4G, I, Supplementary Table 8). Breast tumors enriched with this pericyte-like subset favor CD8^+^ cytotoxic T cells recruitment ^64^, which is coherent with our findings. Based on the genes highly correlating with this subset, pericyte-like presence was associated with enhanced production of growth factors of the FGF family: *FGF7* (Pearson r=0.66, P=1.59E-57), IGF1 (Pearson r=0.62, P=2.58E-48), *FGF2* (Pearson r=0.62, P=2.75E-49) and *IL33* (Pearson r=0.65, P=1.37E-55) (Figure 4H-I, Supplementary Table 8). These genes are important tumor-promoting factors involved in cancer cell proliferation, survival, angiogenesis for the FGF and IGF family members, and inflammation, tissue damage, and fibrosis for *IL33*. In addition to its important role in tissue repair and remodeling, IL33 is also a powerful inducer of T_H_2cell response and might interfere with the establishment of a type-1 helper T (T_H_1) cell anti-tumor response, contributing to the failure of a full CD8^+^ cytotoxic T cell activation ^65^.

Altogether, our data suggest that the microenvironment of HR^+^/HER2^-^ ILC tumors is enriched in iCAFs, which contribute to settle a pro-inflammatory and inappropriate pro-tumoral microenvironment, by maintaining an overall ineffective anti-tumor response.

### Overall coordination of immune and stromal components in HR^+^/HER2^-^ ILCs

To investigate how these actors interact in HR^+^/HER2^-^ ILCs, we performed a correlation matrix of all cell types (Cibersort Absolute Score, CibAbs), CAF subtypes and functional signatures in the whole-transcriptome dataset (446 HR^+^/HER2^-^ ILCs, 2,600 HR^+^/HER2^-^ IDCs). We observed two large clusters along the diagonal of the matrix (Figure 5A).

**Figure 5:**
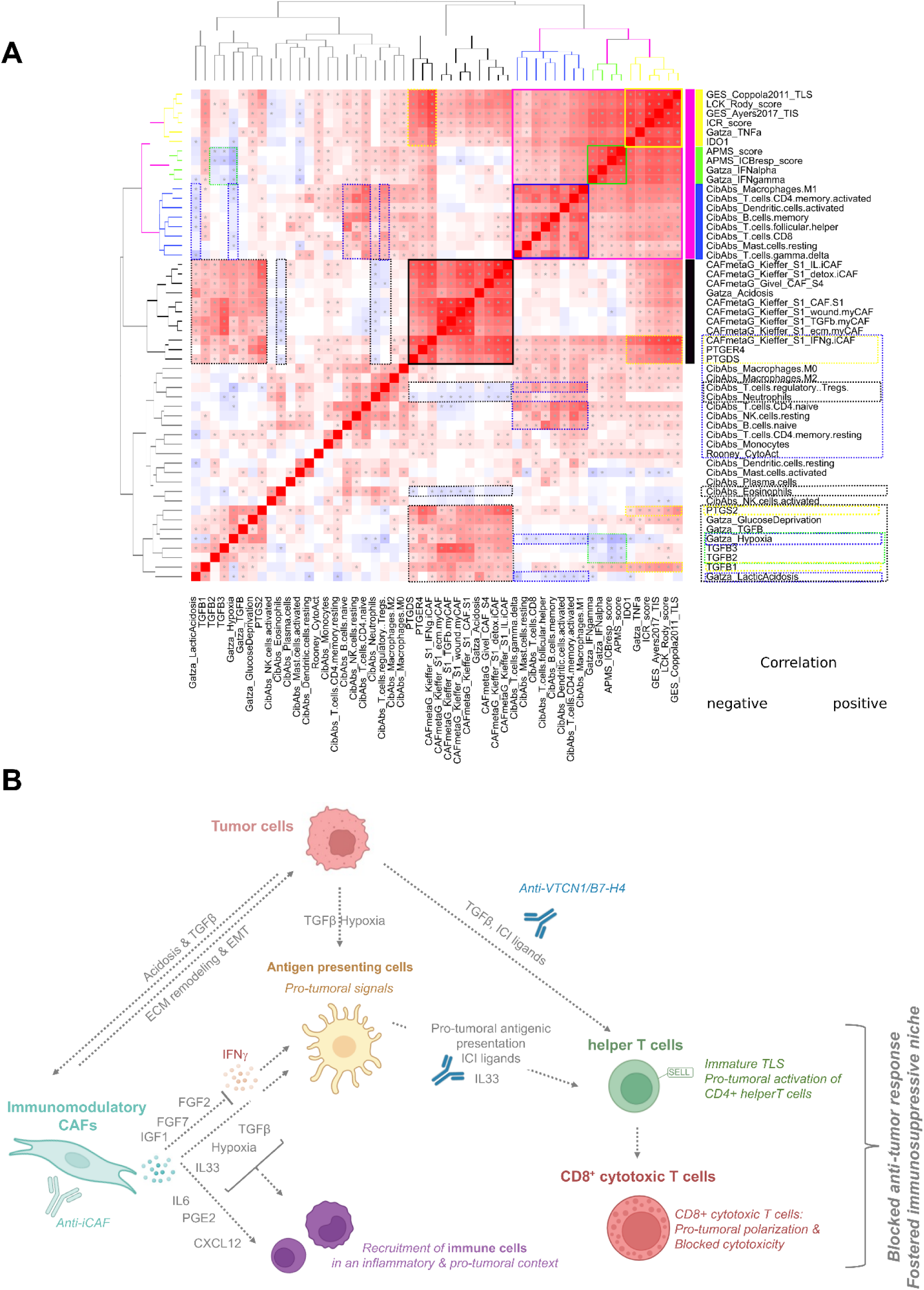
Overall coordination of immune and stromal features in HR^+^/HER2^-^ ILCs. (A) Correlation matrix of score, signatures, and features investigated throughout the study in HR^+^/HER2^-^ ILCs. Solid frame highlighted the strongest correlation and dash frame the secondary one. Colors were for facilitating the description of correlated features. Blue to red colored scale represented negative to positive correlations. (B) Representation of observed HR^+^/HER2^-^ ILCs ecosystem.

The first large cluster (*pink frame*), gathered three sub-clusters: the first sub-cluster (*yellow frame*) consisted of signatures predictive of response to ICIs and *IDO1*; the second sub-cluster (*green frame*) included signatures of anti-tumor activation (antigen presentation to T cells, interferon secretion); and the third sub-cluster (*blue frame*) comprised most of the anti-tumor effectors (type 1 macrophages, cytotoxic T cells, gamma delta T cells) and activated leukocyte states (activated memory helper T cells, activated dendritic cells, …). Altogether, the pink frame showed, as expected, that signatures predictive of response to ICIs, IFN signatures, and anti-tumor response actors were positively correlated. Interestingly, we found that the group of signatures predictive of response to ICIs (*yellow frame*), also strongly correlated with the IFNγ-iCAF metagene and, to a lesser extent with prostaglandin members and *TGFB1* global expression (*dashed yellow frame*). The correlation of anti-tumor immunity actors with known inhibitors of this response suggested a regulatory feedback loop involving *IDO1*, as well as *TGFB1* and *PTGE2* members. Antigen presentation and IFN responses (*green frame*) were anti-correlated with hypoxia, *TGFB2* and *TGFB3* (*dashed green frame*), showing a direct inhibitory link between these actors. Finally, activated immune cells (*blue frame*) showed a significant correlation with their naive counterparts from which they derived, and with regulatory T cells, which likely act as a regulatory loop as well (*dashed blue frame*). Interestingly, they were negatively correlated with hypoxia and lactic acidosis, again suggesting that these factors directly contribute to the inhibition of immune cell activation.

The second large cluster (*black frame*) contained all fibroblast subsets, the signature for acidosis, and members of the prostaglandin family. The sub-cluster of IFNγ-iCAF and prostaglandin members specifically correlated with anti-tumor activation profiles, as previously mentioned (*yellow frames*). Additionally, all fibroblast subsets positively correlated with the pro-tumor cluster composed of *PTGS2*, glucose deprivation, hypoxia, lactic acidosis, and all three TGFβ members. This highlights the link between fibroblast subsets and general pro-tumor and immunosuppressive mechanisms. In addition, fibroblast subsets showed negative correlation with granulocytes (eosinophils and neutrophils), and regulatory T cells. These are also pro-tumor factors whose presence seems to be mutually exclusive with fibroblast subsets.

Altogether, our findings reveal strong coordination between tumor cells and fibroblasts that inhibits anti-tumor cytotoxic CD8^+^ T cell activation by blocking *ad hoc* antigen presentation and secreting immunosuppressive factors (Figure 5B).

## Discussion

In this study, we used multi-omics and multi-scale analysis to explore and compare the immune infiltrates of HR^+^/HER2^-^ ILCs and HR^+^/HER2^-^ IDCs. Single-cell analysis provided higher resolution, while bulk analysis enabled validation at a larger scale; both approaches are therefore complementary. Our single-cell analysis was, however, limited to the only available dataset of ILCs annotated for HR status in the literature. The generation of larger controlled and well-balanced cohorts is required to better profile the immune landscape of BC subtypes at single-cell precision. One strength of our work was the use of a homogeneous cohort of HR^+^/HER2^-^ BC samples, the classical subtype of ILCs, allowing us to limit inter-molecular-subtype heterogeneity.

We first showed that immune infiltration was similar in quantity between the two pathological types but differed in prognosis and quality. TILs infiltration was associated with better prognosis only in HR^+^/HER2^-^IDC samples, but not in HR^+^/HER2^-^ ILCs. However, HR^+^/HER2^-^ ILCs were enriched in CD8^+^ cytotoxic T cells compared to HR^+^/HER2^-^ IDCs. We collected additional data suggesting better immune organization and activation potential of CD8^+^ cytotoxic T cells in HR^+^/HER2^-^ ILCs. Nevertheless, we observed signs of poor and defective anti-tumor activity in HR^+^/HER2^-^ ILCs compared to HR^+^/HER2^-^ IDCs: lower expression of IFNα and IFNγ, defective antigen-presenting cells, increased expression of inhibitory immune checkpoints, low granzyme degranulation, etc. Altogether, our data suggest that although CD8^+^ cytotoxic T cells infiltrate HR^+^/HER2^-^ ILC tumors, they do not achieve an effective anti-tumor response that would indicate a potential benefit from immunotherapy. In this context, they do not display the exhaustion profile that is typical of the complete anti-tumoral response observed in TNBCs. This profile has been associated with better prognosis and ICI benefits ^66^ ^67^. Indeed, despite expressing higher levels of *PD-1*, HR^+^/HER2^-^ ILCs do not overexpress *LAG3, HAVCR2*, or *CD274/PD-L1*, which are important markers of T cell exhaustion following anti-tumor activation ^65^ ^66^. The absence of *CD274/PD-L1* overexpression in HR^+^/HER2^-^ ILC tumors also suggests that using anti-PD-1 or anti-PD-L1 therapies may not be the appropriate strategy to activate CD8^+^ cytotoxic T cells in this tumor subtype.

Several factors may have impeded anti-tumor activity, and we found that some could play a key role in understanding the immune ecosystem of HR^+^/HER2^-^ ILCs and, interestingly, can be targeted. *VTCN1/B7-H4* is one of them. VTCN1/B7-H4 can be expressed by either epithelial tumor cells or antigen-presenting cells. Analysis of single-cell RNA sequencing data revealed that luminal epithelial tumor cells were the primary source of *VTCN1/B7-H4* compared to myeloid cells. Although the presence of its receptor on T cells remains uncertain, VTCN1/B7-H4 has been consistently described as an inhibitory member of the B7 family ^70^ ^71^ ^72^ and is associated with shorter patient survival across multiple cancer types ^73^ ^74^ ^75^. Recent studies have identified B7-H4 as a shared onco-fetal immune tolerance checkpoint, with expression promoted by progesterone in both placental and cancer tissues. This leads to suppression of tumor-associated antigen-specific CD8⁺ cytotoxic T cell immunity and promotes tumor progression ^76^. In breast cancer, VTCN1/B7-H4 is associated to “immune-cold” tumor phenotypes, resistance to immunotherapy, and reduced immune activation of tumor-infiltrating immune cells ^77^ ^78^ ^79,80^ ^81^ ^82^. Notably, VTCN1/B7-H4 has emerged as a promising target for immune checkpoint blockade. Targeting VTCN1/B7-H4 enhances the efficacy of immune checkpoint blockade in preclinical models ^76^, and a VTCN1/B7-H4-directed antibody-drug conjugate (ADC) (BG-C9074) has shown promising preliminary results in a phase I trial in advanced solid tumors, including breast cancer ^83^. Together, these findings highlight VTCN1/B7-H4 as a relevant immunosuppressive mechanism in HR^+^/HER2^-^ ILCs and support its potential as a therapeutic target in this pathological type.

Another central factor appears to be *IL33*, which was identified as one of the most DEG enriched in HR^+^/HER2^-^ ILCs, and reported to be produced by CAFs ^84^ ^65^. IL33, along with other factors produced by iCAFs, has been identified as a key player with pleiotropic functions in inflammatory diseases. This cytokine has an ambiguous role in T cell activation and maturation, which largely depends on the environmental context, but it contributes to inflammation-triggered TLSs in cancer tissues and drives immune tolerance and resistance to cancer therapy ^85^ ^86^ two conditions we observed in HR^+^/HER2^-^ ILC samples. IL33 plays an important role in communication between fibroblasts and immune cells, shaping an overall defective anti-tumor immune response. IL33 also has a role in fibroblast maturation and fibrosis. Specifically, we found a correlation between *IL33* and the IL-iCAF signature in HR^+^/HER2^-^ ILCs. IL33, produced by CAFs, participates in a self-amplifying loop that increases their proliferation, promotes inflammatory cytokine production, and stimulates the secretion of chemokines to attract immune cells. It also increases extracellular matrix synthesis and tumor cell migration, as described in other cancers ^87^ ^88^. This is consistent with the massive stroma observed in ILC tumors, which remains enriched with iCAFs rather than myCAFs. It will be interesting to investigate whether this specific composition of CAF subtypes is somehow related to the Indian chain-like profiles characteristic of ILCs. Further studies are required to investigate in greater detail the role of IL33 in the ecosystem of HR^+^/HER2^-^ ILCs. Interestingly, two anti-IL33 monoclonal antibodies, Itepekimab and Tozorakimab (NCT07484230 and NCT05166889, respectively), are currently tested in clinical trials, and could be of interest for treating HR^+^/HER2^-^ ILCs ILC tumors.

Altogether, our study suggests that the “reactivation” of infiltrating CD8^+^ cytotoxic T cells in HR^+^/HER2^-^ ILC tumors is probably not sufficient to elicit an effective anti-tumor immune response, as these cells suffer from improper activation, at least in part due to an important iCAFs contexture. A strategy relying in activating the anti-tumor immune system should be more appropriate in HR^+^/HER2^-^ ILCs ILCs. Indeed, we show that HR^+^/HER2^-^ ILCs display a complex ecosystem that is distinctly different from that of HR^+^/HER2^-^IDCs, suggesting that targeting the stroma, and notably fibroblasts, and VTCN1/B7-H4 inhibitory signal, is an option that deserves specific attention. Finally, our study highlights the importance of studying this pathological type, which is often set aside in favor of the more prevalent HR^+^/HER2^-^ IDC type.

## Ethics approval and consent to participate

Our study systematically collected the informed patient’s consent to participate, and the ethics and institutional review board were obtained (see Material and Methods section). The study was approved by our institutional review board.

## Availability of data and materials

This study did not generate any new material. For scRNaseq data, all links or accession numbers are listed in the Materials and Methods section. For bulk RNA seq data, all datasets are publicly available, and references are described in Supplementary Table 1.

## Author’s contributions

Conception and design, E.M. and F.B.; methodology, E.M., M.P., F.B.; validation, E.M.; formal analysis, P.F., M.P., A.Gu., G.L., E.M.; resources, L.B., G.L., A.Go., L.M., L.B.; data curation, F.B., M.P., E.M.; writing—original draft preparation, M.P., E.M., F.B.; writing—review and editing, M.P., E.M., F.B., P.F., A.Gu., A.Go., G.L., L.B., L.M.; supervision, E.M. and F.B. All authors have read and agreed to the published version of the manuscript.

## Acknowledgment

The authors wish to thank the IPC/CRCM Experimental Pathology Platform (ICEP) and MICS platform for their technical support. We would like to thank all the donors who have contributed to the study, and the Direction de la Recherche Clinique et Innovation (DRCI).

## Abbreviations

ADC: antibody drug conjugate
APC: antigen presenting cells
BC: breast cancer
CAF: cancer-associated fibroblasts
CMap: connectivity Map
DEG: differentially expressed genes
DFS: disease-free survival
ER: estrogen receptor
FFPE: formalin-fixed paraffin-embedded
HR: hormone receptor
iCAF: inflammatory CAF
ICI: immune checkpoint inhibitor
ICR: immunologic constant of rejection
IDC: invasive ductal carcinoma
ILC: invasive lobular carcinoma
mtpx IF: multiplexed immunofluorescence
myCAF: cancer-associated myofibroblasts
NCS: normalized connectivity score
NK: Natural Killer
OS: overall survival
PBS: phosphate buffer saline
PCA: principal component analysis
PR: progesterone receptor
ROI: region of interest
scRNAseq: single-cell RNA sequencing
TAM2: type 2 tumor associated macrophages
TCR: T cell receptor
T_H_1: type 1 helper T cells
T_H_2: type 2 helper T cells
TIL: tumor-infiltrating lymphocyte
TIS: tumor inflammation signature
TLS: tertiary lymphoid structure
TN: triple negative
UMI: unique molecular identifier

---

**Figure S1:**
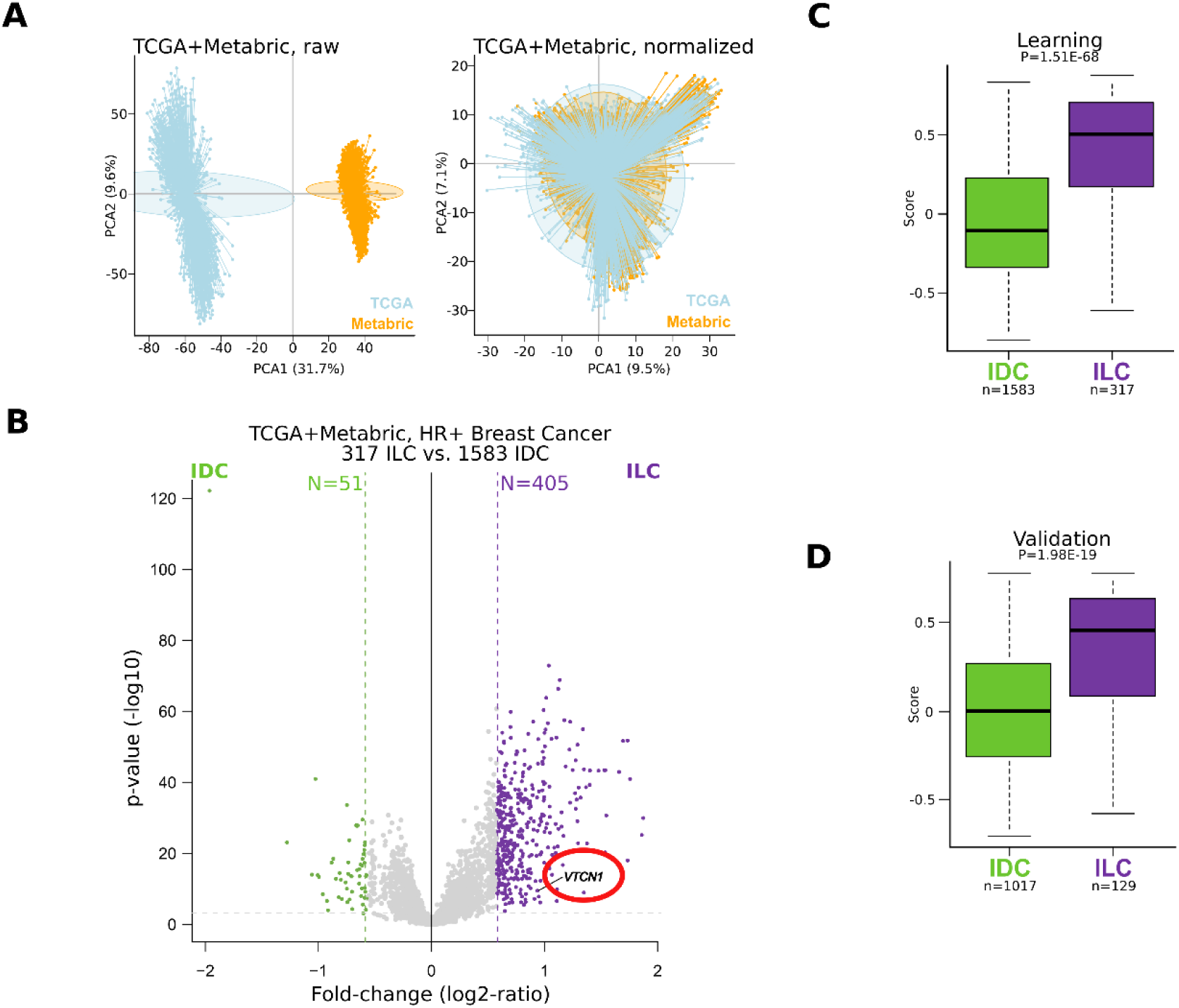
(A) Distribution of TCGA and Metabric raw (left) then normalized (righ) data. PCA; principal component analysis. (B) Volcano plot representing the differential expressed genes in HR^+^/HER2^-^ ILCs *vs* HR^+^/HER2^-^ IDCs from bulk RNA sequencing datasets. VCTN1 is highlighted by a red circle. (C) Robustness of the gene signature to discriminate HR^+^/HER2^-^ ILCs and HR^+^/HER2^-^ IDCs subtypes in the learning dataset. Var; variant, Mod; modality, N; effective, pre; prediction. (D) Robustness of the gene signature to discriminate HR^+^/HER2^-^ ILCs and HR^+^/HER2^-^ IDCs subtypes in the validation dataset. Var; variant, Mod; modality, N; effective, pre; prediction.

**Figure S2:**
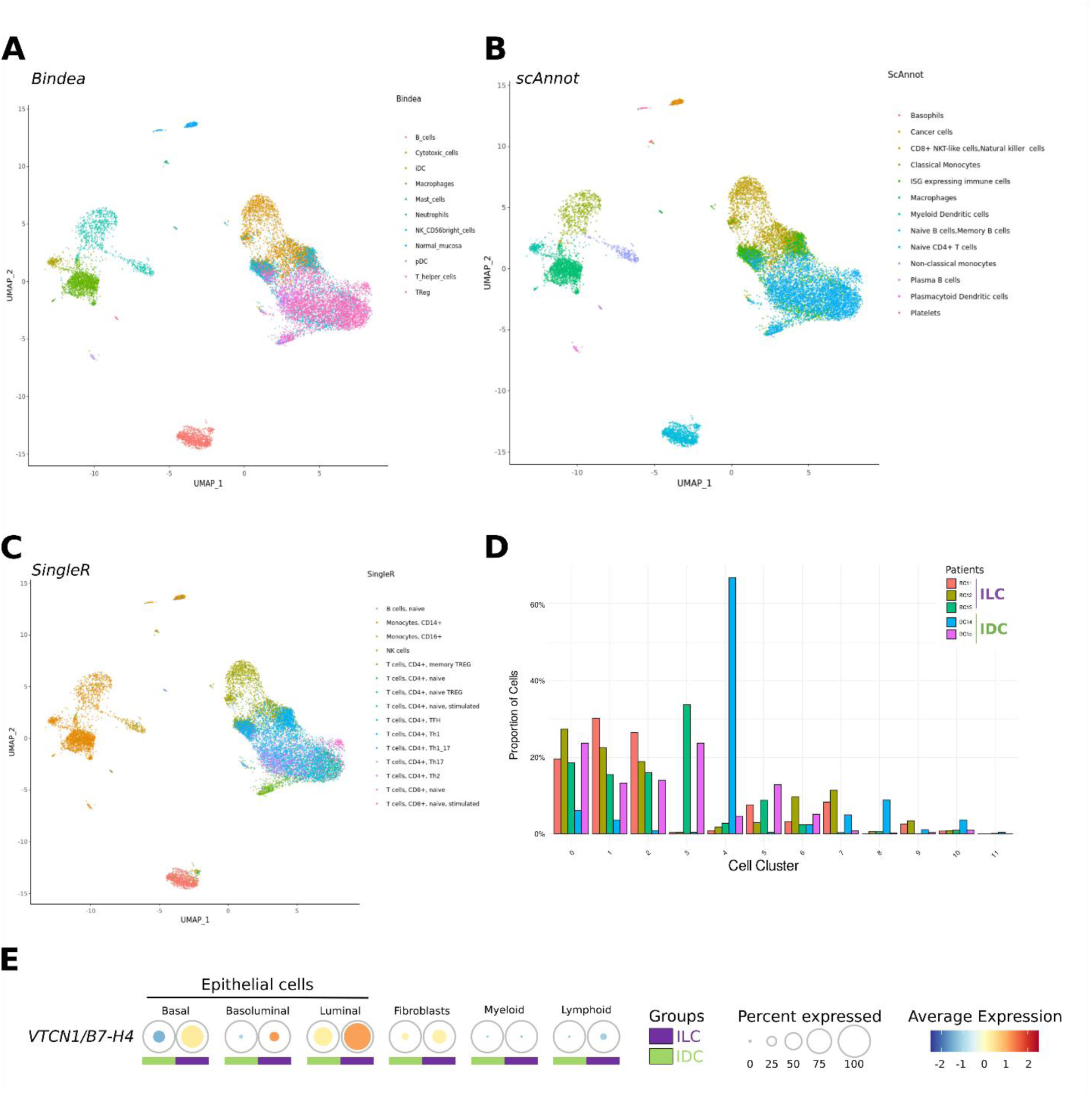
(A) Uniform Manifold Approximation and Projection (uMAP) representing the transfer label from Bindea *et al* immune signatures in the scRNAseq GSE193911 dataset (Onkar *et al*). (B) Uniform Manifold Approximation and Projection (uMAP) representing the transfer label from scAn-not immune signatures in the scRNAseq GSE193911 dataset (Onkar *et al*). (C) Uniform Manifold Approximation and Projection (uMAP) representing the transfer label from Sin-gleR immune signatures in the scRNAseq GSE193911 dataset (Onkar *et al*). (D) Barplot representing the patient contribution for all clusters proportions obtained from the scRNAseq GSE193911 dataset (Onkar *et al*). (E) Dot plots representing the enrichment of the *VTCN1/B7-H4* signature in clusters of the scRNAseq EMTAB10607 dataset (Cords *et al*) in ILCs (*purple*) *vs* IDCs (*light green*). The size of the circle repre-sents the percent of cells expressing the signature and the color represent the absolute strength of the expression

## Bibliography

1. Biglia, N. et al. Increased incidence of lobular breast cancer in women treated with hormone replacement therapy: implications for diagnosis, surgical and medical treatment. Endocr Relat Cancer 14, 549–567 (2007).

2. Dixon, J. M., Renshaw, L., Dixon, J. & Thomas, J. Invasive lobular carcinoma: response to neoadjuvant letrozole therapy. Breast Cancer Res Treat 130, 871–877 (2011).

3. Thomas, M., Kelly, E. D., Abraham, J. & Kruse, M. Invasive lobular breast cancer: A review of pathogenesis, diagnosis, management, and future directions of early stage disease. Semin Oncol 46, 121–132 (2019).

4. Ciriello, G. et al. Comprehensive Molecular Portraits of Invasive Lobular Breast Cancer. Cell 163, 506–519 (2015).

5. Du, T. et al. Invasive lobular and ductal breast carcinoma differ in immune response, protein translation efficiency and metabolism. Sci Rep 8, 7205 (2018).

6. Oesterreich, S., Lucas, P. C., McAuliffe, P. F., Bruno, T. C. & Vignali, D. A. A. Opening the Door for Immune Oncology Studies in Invasive Lobular Breast Cancer. J Natl Cancer Inst 110, 696–698 (2018).

7. Arpino, G., Bardou, V. J., Clark, G. M. & Elledge, R. M. Infiltrating lobular carcinoma of the breast: tumor characteristics and clinical outcome. Breast Cancer Res 6, R149–156 (2004).

8. Dalenc, F. et al. Impact of lobular versus ductal histology on overall survival in metastatic breast cancer: a French retrospective multicentre cohort study. Eur J Cancer 164, 70–79 (2022).

9. Van Baelen, K. et al. Current and future diagnostic and treatment strategies for patients with invasive lobular breast cancer. Ann Oncol 33, 769–785 (2022).

10. Van Baelen, K. et al. Clinical challenges and proposed solutions for patients with invasive lobular breast cancer. Annals of Oncology 36, 1285–1298 (2025).

11. Cardoso, F. et al. Pembrolizumab and chemotherapy in high-risk, early-stage, ER+/HER2- breast cancer: a randomized phase 3 trial. Nat Med 31, 442–448 (2025).

12. Loi, S. et al. Neoadjuvant nivolumab and chemotherapy in early estrogen receptor-positive breast cancer: a randomized phase 3 trial. Nat Med 31, 433–441 (2025).

13. Thompson, E. D. et al. PD-L1 expression and the immune microenvironment in primary invasive lobular carcinomas of the breast. Modern Pathology 30, 1551–1560 (2017).

14. Michaut, M. et al. Integration of genomic, transcriptomic and proteomic data identifies two biologically distinct subtypes of invasive lobular breast cancer. Sci Rep 6, 18517 (2016).

15. Rugo, H. S. et al. Safety and Antitumor Activity of Pembrolizumab in Patients with Estrogen Receptor-Positive/Human Epidermal Growth Factor Receptor 2-Negative Advanced Breast Cancer. Clin Cancer Res 24, 2804–2811 (2018).

16. Voorwerk, L. et al. PD-L1 blockade in combination with carboplatin as immune induction in metastatic lobular breast cancer: the GELATO trial. Nat Cancer 4, 535–549 (2023).

17. Desmedt, C. et al. Immune Infiltration in Invasive Lobular Breast Cancer. JNCI: Journal of the National Cancer Institute 110, 768–776 (2018).

18. Onkar, S. et al. Immune landscape in invasive ductal and lobular breast cancer reveals a divergent macrophage-driven microenvironment. Nat Cancer 4, 516–534 (2023).

19. Bertucci, F. et al. The immunologic constant of rejection classification refines the prognostic value of conventional prognostic signatures in breast cancer. Br J Cancer 119, 1383–1391 (2018).

20. Sabatier, R. et al. Down-regulation of ECRG4, a candidate tumor suppressor gene, in human breast cancer. PLoS One 6, e27656 (2011).

21. Bertucci, F. et al. PDL1 expression is a poor-prognosis factor in soft-tissue sarcomas. Oncoimmunology 6, e1278100 (2017).

22. Lehmann, B. D. et al. Identification of human triple-negative breast cancer subtypes and preclinical models for selection of targeted therapies. J Clin Invest 121, 2750–2767 (2011).

23. Parker, J. S. et al. Supervised risk predictor of breast cancer based on intrinsic subtypes. J Clin Oncol 27, 1160–1167 (2009).

24. Paik, S. et al. A multigene assay to predict recurrence of tamoxifen-treated, node-negative breast cancer. N Engl J Med 351, 2817–2826 (2004).

25. van ’t Veer, L. J., et al. Gene expression profiling predicts clinical outcome of breast cancer. Nature 415, 530–536 (2002).

26. Truntzer, C., Isambert, N., Arnould, L., Ladoire, S. & Ghiringhelli, F. Prognostic value of transcriptomic determination of tumour-infiltrating lymphocytes in localised breast cancer. Eur J Cancer 120, 97–106 (2019).

27. Bindea, G. et al. Spatiotemporal dynamics of intratumoral immune cells reveal the immune landscape in human cancer. Immunity 39, 782–795 (2013).

28. Newman, A. M. et al. Determining cell type abundance and expression from bulk tissues with digital cytometry. Nat Biotechnol 37, 773–782 (2019).

29. Coppola, D. et al. Unique ectopic lymph node-like structures present in human primary colorectal carcinoma are identified by immune gene array profiling. Am J Pathol 179, 37–45 (2011).

30. Ayers, M. et al. Gene expression profiles predict complete pathologic response to neoadjuvant paclitaxel and fluorouracil, doxorubicin, and cyclophosphamide chemotherapy in breast cancer. J Clin Oncol 22, 2284–2293 (2004).

31. Rooney, M. S., Shukla, S. A., Wu, C. J., Getz, G. & Hacohen, N. Molecular and genetic properties of tumors associated with local immune cytolytic activity. Cell 160, 48–61 (2015).

32. Rody, A. et al. T-cell metagene predicts a favorable prognosis in estrogen receptor-negative and HER2-positive breast cancers. Breast Cancer Res 11, R15 (2009).

33. Gatza, M. L. et al. A pathway-based classification of human breast cancer. Proc Natl Acad Sci U S A 107, 6994–6999 (2010).

34. Benjamini, Y. & Hochberg, Y. Controlling the False Discovery Rate: A Practical and Powerful Approach to Multiple Testing. Journal of the Royal Statistical Society Series B: Statistical Methodology 57, 289–300 (1995).

35. Cords, L. et al. Cancer-associated fibroblast classification in single-cell and spatial proteomics data. Nat Commun 14, 4294 (2023).

36. Windhager, J. et al. An end-to-end workflow for multiplexed image processing and analysis. Nat Protoc 18, 3565–3613 (2023).

37. Greenwald, N. F. et al. Whole-cell segmentation of tissue images with human-level performance using large-scale data annotation and deep learning. Nat Biotechnol 40, 555–565 (2022).

38. Oesterreich, S. et al. Clinicopathological Features and Outcomes Comparing Patients With Invasive Ductal and Lobular Breast Cancer. J Natl Cancer Inst 114, 1511–1522 (2022).

39. Bleiweiss IJ. Pathology of breast cancer. In: UpToDate. Chagpar AB, Vora SR (eds.). Waltham, MA: UpToDate, 2024. in.

40. Peyraud, F. et al. Tertiary lymphoid structures and cancer immunotherapy: From bench to bedside. Med 6, 100546 (2025).

41. Ayers, M. et al. IFN-γ-related mRNA profile predicts clinical response to PD-1 blockade. J Clin Invest 127, 2930–2940 (2017).

42. Uleri, V. et al. LCK deficiency in CD8 T cells leads to reduced proliferation and increased effector T-cell formation in mice. Preprint at 10.1101/2025.08.28.672860 (2025).

43. Gatza, M. L. et al. Analysis of tumor environmental response and oncogenic pathway activation identifies distinct basal and luminal features in HER2-related breast tumor subtypes. Breast Cancer Res 13, R62 (2011).

44. Simon, S. & Labarriere, N. PD-1 expression on tumor-specific T cells: Friend or foe for immunotherapy? Oncoimmunology 7, e1364828 (2017).

45. Whelan, S. et al. PVRIG and PVRL2 Are Induced in Cancer and Inhibit CD8+ T-cell Function. Cancer Immunology Research 7, 257–268 (2019).

46. Scheffges, C. et al. Identification of CD160-TM as a tumor target on triple negative breast cancers: possible therapeutic applications. Breast Cancer Res 26, 28 (2024).

47. Martin, A. S. et al. VISTA expression and patient selection for immune-based anticancer therapy. Front. Immunol. 14, 1086102 (2023).

48. Murphy, K. M., Nelson, C. A. & Šedý, J. R. Balancing co-stimulation and inhibition with BTLA and HVEM. Nat Rev Immunol 6, 671–681 (2006).

49. Ren, S. et al. The immune checkpoint, HVEM may contribute to immune escape in non-small cell lung cancer lacking PD-L1 expression. Lung Cancer 125, 115–120 (2018).

50. Zhou, L. et al. B7H4 expression in tumor cells impairs CD8 T cell responses and tumor immunity. Cancer Immunol Immunother 69, 163–174 (2020).

51. Wang, J.-Y. & Wang, W.-P. B7-H4, a promising target for immunotherapy. Cellular Immunology 347, 104008 (2020).

52. Murphy, K. M., Nelson, C. A. & Šedý, J. R. Balancing co-stimulation and inhibition with BTLA and HVEM. Nat Rev Immunol 6, 671–681 (2006).

53. Musa, M. Single-cell analysis on stromal fibroblasts in the microenvironment of solid tumours. Advances in Medical Sciences 65, 163–169 (2020).

54. Gandellini, P. et al. Complexity in the tumour microenvironment: Cancer associated fibroblast gene expression patterns identify both common and unique features of tumour-stroma crosstalk across cancer types. Seminars in Cancer Biology 35, 96–106 (2015).

55. Croizer, H. et al. Deciphering the spatial landscape and plasticity of immunosuppressive fibroblasts in breast cancer. Nat Commun 15, 2806 (2024).

56. Licaj, M. et al. Residual ANTXR1+ myofibroblasts after chemotherapy inhibit anti-tumor immunity via YAP1 signaling pathway. Nat Commun 15, 1312 (2024).

57. Biffi, G. & Tuveson, D. A. Diversity and Biology of Cancer-Associated Fibroblasts. Physiol Rev 101, 147–176 (2021).

58. Givel, A.-M. et al. miR200-regulated CXCL12β promotes fibroblast heterogeneity and immunosuppression in ovarian cancers. Nat Commun 9, 1056 (2018).

59. Kieffer, Y. et al. Single-Cell Analysis Reveals Fibroblast Clusters Linked to Immunotherapy Resistance in Cancer. Cancer Discov 10, 1330–1351 (2020).

60. Mhaidly, R. & Mechta-Grigoriou, F. Fibroblast heterogeneity in tumor micro-environment: Role in immunosuppression and new therapies. Semin Immunol 48, 101417 (2020).

61. Thiery, J. Modulation of the antitumor immune response by cancer-associated fibroblasts: mechanisms and targeting strategies to hamper their immunosuppressive functions. Explor Target Antitumor Ther 3, 598–629 (2022).

62. Mhaidly, R. & Mechta-Grigoriou, F. Role of cancer-associated fibroblast subpopulations in immune infiltration, as a new means of treatment in cancer. Immunol Rev 302, 259–272 (2021).

63. Barrett, R. L. & Puré, E. Cancer-associated fibroblasts and their influence on tumor immunity and immunotherapy. Elife 9, e57243 (2020).

64. Costa, A. et al. Fibroblast Heterogeneity and Immunosuppressive Environment in Human Breast Cancer. Cancer Cell 33, 463–479.e10 (2018).

65. Yeoh, W. J., Vu, V. P. & Krebs, P. IL-33 biology in cancer: An update and future perspectives. Cytokine 157, 155961 (2022).

66. Sabatier, R. et al. Prognostic and predictive value of PDL1 expression in breast cancer. Oncotarget 6, 5449–5464 (2015).

67. Jacquemier, J. et al. High expression of indoleamine 2,3-dioxygenase in the tumour is associated with medullary features and favourable outcome in basal-like breast carcinoma. Int J Cancer 130, 96–104 (2012).

68. Sabatier, R. et al. A gene expression signature identifies two prognostic subgroups of basal breast cancer. Breast Cancer Res Treat 126, 407–420 (2011).

69. Bertucci, F. et al. Gene expression profiling shows medullary breast cancer is a subgroup of basal breast cancers. Cancer Res 66, 4636–4644 (2006).

70. Sica, G. L. et al. B7-H4, a molecule of the B7 family, negatively regulates T cell immunity. Immunity 18, 849–861 (2003).

71. Prasad, D. V. R., Richards, S., Mai, X. M. & Dong, C. B7S1, a novel B7 family member that negatively regulates T cell activation. Immunity 18, 863–873 (2003).

72. Zang, X. et al. B7x: a widely expressed B7 family member that inhibits T cell activation. Proc Natl Acad Sci U S A 100, 10388–10392 (2003).

73. Kryczek, I. et al. Relationship between B7-H4, regulatory T cells, and patient outcome in human ovarian carcinoma. Cancer Res 67, 8900–8905 (2007).

74. Podojil, J. R. & Miller, S. D. Potential targeting of B7-H4 for the treatment of cancer. Immunol Rev 276, 40–51 (2017).

75. Smith, J. B., Stashwick, C. & Powell, D. J. B7-H4 as a potential target for immunotherapy for gynecologic cancers: a closer look. Gynecol Oncol 134, 181–189 (2014).

76. Yu, J. et al. Progestogen-driven B7-H4 contributes to onco-fetal immune tolerance. Cell 187, 4713–4732.e19 (2024).

77. Wescott, E. C. et al. Epithelial Expressed B7-H4 Drives Differential Immunotherapy Response in Murine and Human Breast Cancer. Cancer Res Commun 4, 1120–1134 (2024).

78. Gruosso, T. et al. Spatially distinct tumor immune microenvironments stratify triple-negative breast cancers. J Clin Invest 129, 1785–1800 (2019).

79. Kim, N. I., Park, M. H., Kweon, S.-S. & Lee, J. S. B7-H3 and B7-H4 Expression in Breast Cancer and Their Association with Clinicopathological Variables and T Cell Infiltration. Pathobiology 87, 179–192 (2020).

80. Wang, L., Yang, C., Liu, X.-B., Wang, L. & Kang, F.-B. B7-H4 overexpression contributes to poor prognosis and drug-resistance in triple-negative breast cancer. Cancer Cell Int 18, 100 (2018).

81. Song, X. et al. Pharmacologic Suppression of B7-H4 Glycosylation Restores Antitumor Immunity in Immune-Cold Breast Cancers. Cancer Discov 10, 1872–1893 (2020).

82. Sousa, L. G. et al. Spatial Immunoprofiling of Adenoid Cystic Carcinoma Reveals B7-H4 Is a Therapeutic Target for Aggressive Tumors. Clin Cancer Res 29, 3162–3171 (2023).

83. Perez, C. A. et al. First-in-human study of BG-C9074, a B7-H4-targeting ADC in patients with advanced solid tumors: Preliminary results of the dose-escalation phase. JCO 43, 3033–3033 (2025).

84. Donahue, K. L. et al. Oncogenic KRAS-Dependent Stromal Interleukin-33 Directs the Pancreatic Microenvironment to Promote Tumor Growth. Cancer Discovery 14, 1964–1989 (2024).

85. Amisaki, M. et al. IL-33-activated ILC2s induce tertiary lymphoid structures in pancreatic cancer. Nature 638, 1076–1084 (2025).

86. Lamorte, S. et al. Lymph node macrophages drive immune tolerance and resistance to cancer therapy by induction of the immune-regulatory cytokine IL-33. Cancer Cell 43, 955–969.e10 (2025).

87. Eguchi, S. et al. IL-33 released during liver resection facilitates intrahepatic cholangiocarcinoma growth via cytokine secretion in cancer-associated fibroblasts. Br J Cancer 134, 519–529 (2026).

88. Yang, L., Zhou, J., Chen, S., Wang, Y. & Huang, S. CAF derived IL-33 mediated EMT to promote the metastasis of LSCC cells. Eur J Med Res 30, 1147 (2025).

